# IL-10 Targets IRFs to Suppress IFN and Inflammatory Response Genes by Epigenetic Mechanisms

**DOI:** 10.1101/2024.11.07.622491

**Authors:** Bikash Mishra, Mahesh Bachu, Claire Wingert, Ruoxi Yuan, Vidyanath Chaudhary, Caroline Brauner, Richard Bell, Lionel B. Ivashkiv

## Abstract

Interleukin-10 (IL-10) is pivotal in suppressing inflammation and innate immune activation, in large part by suppressing induction of genes by potent inflammatory factors such as TLR ligands. Despite decades of research, molecular mechanisms underlying this inhibition have not been resolved. This study utilized an integrated epigenomic analysis of gene transcription, chromatin accessibility, histone modifications and transcription factor binding to investigate IL-10-mediated suppression of LPS and TNF responses in primary human monocytes. Instead of inhibiting core TLR4-activated pathways such as NF-κB, MAPK-AP-1 and TBK1-IRF3 signaling, IL-10 targeted IRF transcription factor activity and DNA binding, particularly IRF5 and an IRF1-mediated amplification loop that is operative in monocytes. This resulted in suppression of inflammatory NF-κB target genes, in whose activation IRFs play an amplifying role, and near-complete suppression of interferon-stimulated genes. Mechanisms of TLR4 and TNFR target gene inhibition included downregulation of chromatin accessibility, de novo enhancer formation, and IRF1-associated H3K27ac activating histone marks. These results provide a mechanism by which IL-10 suppresses inflammatory NF-κB target genes, highlight the role of IRF1 in inflammatory gene expression, and describe an underappreciated suppression of IFN responses by epigenetic mechanisms.

## Introduction

IL-10 is a key suppressor of innate immune activation and inflammatory responses ^1, 2, 3, 4^. A well-recognized mechanism of IL-10 action is suppression of production of inflammatory cytokines encoded by NF-κB target genes such as *TNF*, *IL6*, *IL1B* and *IL12B* by myeloid cells including monocytes, dendritic cells, and macrophages ^4, 5^. The absence of IL-10 results in hyperactivation of innate immune and inflammatory responses upon environmental challenges, and the spontaneous development of inflammatory conditions, most notably colitis in IL-10 deficient mice ^6^. The importance of monocyte and macrophage suppression in preventing excessive inflammation and associated diseases has been demonstrated by myeloid cell-specific deletion of the IL-10 receptor or of the major effector of IL-10 signaling STAT3 ^7, 8^. Mutations in STAT3 or the IL-10 receptor result in inflammatory bowel diseases in humans ^6, 7, 8, 9, 10^. The importance of IL-10 as a negative regulator of innate immunity and inflammation have led to extensive efforts to understand signaling pathways activated by IL-10 and mechanisms by which it suppresses inflammatory gene activation ^11^.

Upon binding to its receptor composed of IL-10R1 and IL-10R2 subunits, IL-10 initiates a core signaling pathway involving activation of tyrosine protein kinases JAK1 and TYK2, and transcription factor STAT3. STAT3 is required for the transcriptional response to IL-10, and for the core IL-10 function of suppressing induction of inflammatory genes by microbial products, DAMPs and cytokines ^4, 12, 13^. However, it has proved challenging to identify mechanisms by which IL-10-STAT3 signaling suppresses inflammatory gene induction ^14^. STAT3 does not bind to and directly suppress inflammatory genes ^15, 16^, and an early hypothesis proposed in the 1980s that IL-10-STAT3 signaling suppresses TLR-induced activation of NF-κB or MAPK signaling was not supported by a large body of data ^1, 5, 11, 17, 18^. This led to the alternative idea that IL-10-STAT3 signaling inhibits inflammatory genes indirectly, via induction of transcriptional repressors that then bind to and inhibit inflammatory genes ^11^. Although various IL-10-induced transcriptional repressors have been identified including BATF3, BCL3, NFIL3, and SBNO2, none of the identified factors played a key nonredundant role in suppressing core inflammatory genes such as *TNF* and *IL12B* ^5, 19, 20, 21^. More recently, another line of investigation showed that IL-10 counters the metabolic shift triggered by inflammatory cues in macrophages, partly by regulating glucose uptake and glycolysis while favoring oxidative phosphorylation and inhibiting mTOR signaling ^22^. The IL-10-induced metabolic changes explained suppression of inflammasome activation, but mechanisms by which IL-10 suppresses inflammatory gene transcription remained unresolved. Metabolic reprogramming can affect chromatin states by modulating levels of substrates, cofactors, and energy utilized by epigenetic remodeling enzymes. However, to date there is limited evidence that IL-10 suppresses inflammatory genes by epigenetic mechanisms ^16, 23, 24^.

The inhibitory effects of IL-10 on inflammatory gene induction are typically studied in mouse macrophages stimulated with prototypical activators such as TLR4 ligand LPS. TLR4 activates a pro-inflammatory signaling branch via adaptor molecule MyD88 that activates NF-κB and MAPKs that drive expression of inflammatory genes such as *TNF*, *IL6* and *IL1B*. IRF5, which is activated via MyD88 by a unknown mechanism ^25^ contributes to and amplifies inflammatory gene expression, at least in part by cooperation with NF-κB ^26, 27^. TLR4 also activates a Trif-mediated signaling pathway that contributes to inflammatory gene expression but whose predominant function is induction of interferon-stimulated genes (ISGs). ISGs are induced indirectly via activation of production of IFN-b, which activates Jak-STAT signaling and transcription factor complex ISGF3 (comprised of STAT1, STAT2 and IRF9) which binds to ISRE DNA elements at ISG genomic loci ^28, 29^. Induction of *IFNB*, and thus full activation of ISGs, is typically completely dependent on Trif-mediated activation of IRF3. In contrast to IRF5, IRF1, whose expression is induced by IFN-γ and type I IFNs, has been mainly implicated in induction of ISGs via binding to genomic elements that closely resemble the ISRE ^28, 29, 30, 31^. However, IRF1 was implicated in induction of *IFNB* in the paper that first reported its molecular cloning ^32, 33, 34^ and more recent work has shown IRF1 binding at inflammatory gene loci ^35^, suggesting a potentially broader function for this transcription factor. As IL-10 has not been appreciated to suppress IFN responses, the regulation of IRFs by IL-10 has not been well investigated.

We reasoned that applying high dimensional genome-wide approaches could help solve the unresolved question of how IL-10 inhibits inflammatory gene induction. Thus, we utilized an integrated epigenomic approach combining RNAseq, ATACseq and CUT&RUN to investigate regulation of LPS and TNF responses by IL-10 in primary human monocytes. Monocytes are key producers of inflammatory cytokines whose activation needs to be regulated to maintain effective host defense while preventing collateral tissue damage or cytokine storm. Freshly isolated monocytes correspond directly to cells that exit the circulation and enter inflamed tissues, and thus analysis of these cells reflects a high degree of biological relevance as it models in vivo activation. We found that in human monocytes IRF1 and IRF5 play a key role in *IFNB*, ISG and inflammatory gene induction, and that IL-10 suppresses TLR4- and TNFR-induced gene expression by suppressing IRFs rather than inhibiting core NF-κB, MAPK-AP-1, or TBK1-IRF3 signaling pathways. IL-10 utilizes epigenetic mechanisms involving chromatin accessibility, enhancer formation, and IRF1-associated histone modification to suppresses IRF-dependent interferon responses more strongly than canonical inflammatory NF-κB target genes. Our study highlights inhibition of IFN responses as a biological activity of IL-10 and unveils a previously unrecognized mechanism of IL-10-mediated suppression of TLR4- and TNFR-induced gene expression.

## Results

### IL-10 preferentially suppresses IFN relative to inflammatory responses in LPS- and TNF-stimulated monocytes

We used RNAseq to analyze the effects of IL-10 on the TLR4-induced gene expression profile in primary human monocytes (Fig. 1a, Extended Data Fig. 1a-b). As expected, LPS activated inflammatory and IFN pathways and induced expression (FDR < 0.05, fold change > 2) of known pro-inflammatory NF-κB target genes such as *IL6* and *TNF*, and interferon response genes like *CXCL10* and *ISG15*, which was confirmed by qPCR for multiple independent donors (Fig. 1b). Analysis of the effects of IL-10 on the LPS-induced response using Gene Set Enrichment Analysis (GSEA) confirmed the expected suppression of inflammatory genes by IL-10 (Fig. 1c, top panel). Surprisingly, LPS-induced IFN response genes were more significantly and consistently suppressed by IL-10 than canonical inflammatory genes (Fig. 1c, bottom panel); inhibition of ISGs by IL-10 was confirmed by qPCR in multiple donors (Fig. 1b). To identify the full spectrum of LPS-inducible genes that were suppressed by IL-10, we initially clustered genes based on pattern of expression and performed pathway analysis (Fig. 1d-e). Strikingly, gene cluster 2 whose induction was strongly suppressed by IL-10 was most significantly enriched for IFN response genes. In contrast, LPS-induced cluster 3 genes that were only moderately suppressed by IL-10 were most significantly enriched for inflammatory NF-κB pathways (Fig. 1e). In a complementary approach, we identified LPS-induced genes suppressed by at least 40% by IL-10 (436 out of 1745 genes, Extended Data Fig. 1c-d). The overwhelming majority (92.4%) of these genes fell into clusters 2 or 3 (Extended Data Fig. 1e); this convergence of results provides a core gene set of IL-10-suppressed genes. Further analysis of the core set of IL-10 suppressed genes (40% or more suppressed) showed most significant enrichment of IFN pathways, with less significant enrichment of inflammatory and NF-κB pathways (Fig. 1f). Heatmaps depicting expression of Hallmark pathway inflammatory and IFN response genes clearly show that ISGs were more strongly suppressed than inflammatory genes (Fig. 1g-h). In accord with previous literature ^14, 17, 19^, we found that IL-10 did not inhibit LPS-mediated activation of NF-κB signaling (Extended Data Fig. 1f). These results extend previously reported IL-10-mediated suppression of inflammatory genes to primary human monocytes and reveal an unexpected strong suppression of the LPS-induced IFN response.

**Figure 1.**
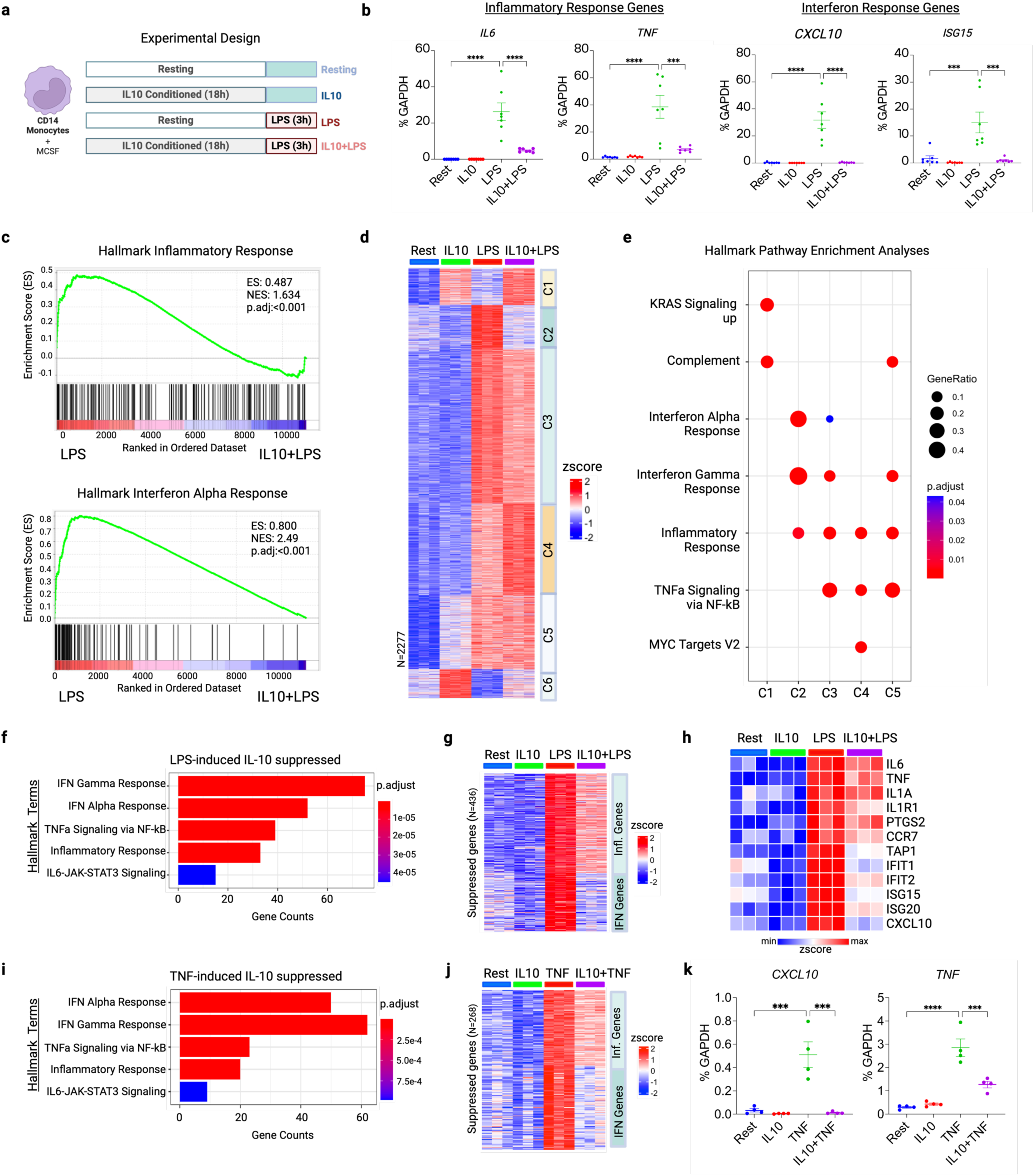
Preferential suppression of IFN relative to inflammatory responses in LPS- and TNF-stimulated monocytes by IL-10. **a,** Experimental design. Human monocytes were cultured –/+ IL-10 (100 ng/mL) for 18 hours and then stimulated with LPS (10 ng/ml) for 3 hours. **b,** mRNA of indicated genes was measured by qPCR and normalized relative to *GAPDH* mRNA in cells stimulated as indicated (dots correspond to independent donors; n = 7). **c-j,** analysis of RNAseq data obtained using monocytes from 3 independent donors and treated as depicted in **a** **c,** GSEA of RNA-seq data performed on DEGs ranked by log2CPMs in LPS and IL-10+LPS treated human monocytes. The analysis utilized Hallmark gene sets from Broad Institute and LPS-regulated IFN and inflammatory pathways are displayed. (NES: Normalized Expression Score, ES: Expression Score) **d,** k-means clustering analysis (k=6) conducted on differentially upregulated genes in any pairwise comparison relative to resting control. **e-f,** Hallmark pathway enrichment analyses performed **e,** on clusters identified in Fig. 1d, f, on LPS-induced genes identified to be suppressed genes by IL-10. **g-h,** Heatmaps of **g,** LPS-induced genes identified to be suppressed genes by IL-10, clustered by Inflammatory and IFN response genes, **h,** representative IL-10-suppressed inflammatory and IFN response genes induced by LPS. **i,** Hallmark pathway enrichment analysis performed on TNF-induced genes identified to be suppressed genes by IL-10. **j,** Heatmap of TNF-induced genes identified to be suppressed genes by IL-10, clustered by Inflammatory and IFN response genes. **k,** mRNA of *CXCL10* and *TNF* was measured by qPCR and normalized relative to *GAPDH* mRNA in cells stimulated as indicated (n=4 independent donors). For d,g,h, and j, z-score normalized data was used to make heatmaps. For b and k, data are depicted as mean ± SEM. ***p < 0.0005; ****p < 0.0001 by One-way ANOVA with Tukey’s multiple comparisons test.

TNF induces expression of inflammatory and IFN response genes but utilizes distinct upstream signaling pathways, most notably induction of the IFN response is dependent on IRF1 rather than IRF3 ^36, 37, 38^. We next performed similar RNAseq experiments to determine whether IL-10 inhibits TNF-induced inflammatory and IFN response genes in a similar manner as inhibition of LPS (Extended Data Fig. 1g-h). IL-10 strongly suppressed TNF-induced interferon genes, but only moderately suppressed inflammatory NF-κB target genes (Fig. 1i-k and Extended Data Fig. 1i-l). Collectively, these results reveal strong inhibition of IFN responses in human monocytes as a biological function of IL-10.

### IL-10 suppresses H3K27ac at genomic regions enriched in IRF1 binding motifs

The similarity of suppression of LPS and TNF responses, the gene-specific nature of suppression, the resistance of subsets of LPS-inducible genes to IL-10-mediated inhibition (Fig. 1d, clusters 4 and 5), and the literature showing that IL-10 does not inhibit NF-κB and MAPK signaling ^17^ suggest that IL-10 may inhibit gene expression by an epigenetic chromatin-mediated mechanism. We first tested this notion by using CUT&RUN to obtain the genome-wide profile of histone 3 lysine 27 acetylation (H3K27ac), which marks active gene-regulatory elements (enhancers and promoters) (Extended Data Fig. 2a). LPS significantly increased H3K27ac at 5,749 genomic regions using criteria of FDR ≤ 0.05 and 2-fold (Extended Data Fig. 2b). Of these 5,749 H3K27ac peaks, 3450 (60%) were not called as significant peaks in the IL-10 + LPS condition, (Fig. 2a, termed Group i or ‘LPS unique’, which corresponds to IL-10 suppressed peaks). In contrast, 2299 of the LPS-inducible peaks called were present in the IL-10 + LPS condition; these are termed Group ii and are resistant to IL-10 inhibition. Quantitation of peak intensity using deeptools and visualization on heat maps (Fig. 2b) or violin plots (Fig. 2c) showed that IL-10 highly significantly suppressed Group i H3K27ac peaks to near baseline conditions. Interestingly, IL-10 also significantly suppressed H3K27ac at Group ii peaks, but this suppression was partial and mean peak intensity remained elevated relative to the baseline observed in unstimulated monocytes (Fig. 2b-c). Induced peaks were distributed in intergenic, intronic and promoter regions (Fig. 2d), suggesting that IL-10 targets both enhancer and promoter regulatory elements to suppress associated genes. Representative gene tracks showing suppression of H3K27ac at Group i enhancer and promoter peaks, with preservation of H3K27ac at Group ii peaks are shown in Fig. 2e. In accord with regulation of promoter activity, CUT&RUN analysis of H3K4me3, a positive promoter mark that promotes transcription, showed that genes with decreased transcription and H3K27ac also had decreased H3K4me3, whereas H3K4me3 remained unchanged at gene loci where H3K27ac was not suppressed (Fig. 2e-f). Genes associated with Group i H3K27ac peaks were strongly enriched for hallmark interferon pathways (Extended Data Fig. 2c), and a large fraction of IL-10-supressed genes defined in Fig. 1 were associated with Group i IL-10-suppressed peaks (Extended Data Fig. 2d). Collectively, these results show that IL-10 suppresses LPS-induced activation of chromatin and suggest that this suppression is associated decreased expression of LPS-induced genes.

**Figure 2.**
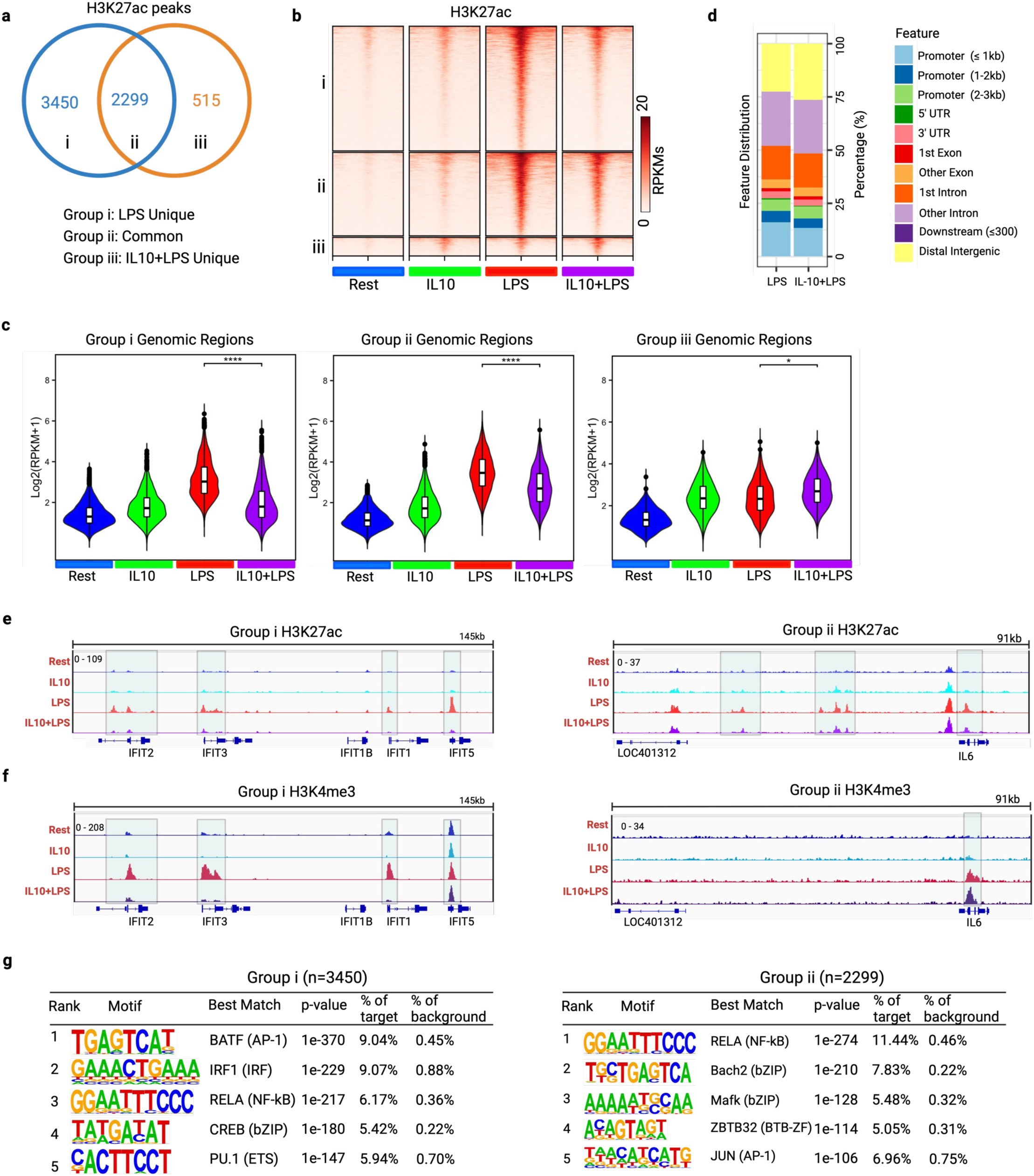
IL-10 suppresses H3K27ac at genomic regions enriched in IRF1 binding motifs. **a-h**, analysis of H3K27ac CUT and RUN data obtained using monocytes from 2 independent donors and treated as depicted in **1a a,** Venn diagram showing numbers of differentially upregulated H3K27ac peaks in LPS and IL-10+LPS treatment relative to resting controls in primary human monocytes. **b,** Heatmap of the normalized signal density of H3K27ac binding at Groups i, ii, and iii from **a** under indicated conditions. The results are presented in RPKM (reads per kilobase per million mapped reads) values within a range of ± 2.0kb around peak centers. **c,** Violin plots showing normalized average signal density of H3K27ac binding at Group-i, Group-ii, and Group-iii peaks shown in **b** under indicated conditions. Data plotted as log2 (RPKM+1) counts of H3K27ac reads. *p < 0.05; ****p < 0.0001 by Two-way ANOVA with Tukey’s multiple comparisons test **d,** Genomic feature distribution of H3K27ac peak coordinates in LPS and IL-10+LPS treated primary human monocytes. **e-g,** Representative Interactive Genome Viewer (IGV) gene tracks of **e,** H3K27ac binding for Group-i (*IFIT* locus, left panel) and Group-ii (*IL6*, right panel), **f,** of H3K4me3 binding for suppressed gene (*IFIT* loci), and **g,** of H3K4me3 binding for non-suppressed gene (*IL6*). H3K4Me3 CUT and RUN data were obtained using monocytes from 2 independent donors. **h,** *De novo* motif analysis results using HOMER on Group-i and Group-ii H3K27ac peaks.

To gain insight into whether Group i and Group ii peaks could be distinguished based on regulation by different transcription factors (TFs), we used HOMER to perform *de novo* motif analysis to identify TF binding sequences enriched under H3K27ac peaks. Both Group i and Group ii peaks were enriched for AP-1 and NF-κB motifs, whereas only Group i peaks, which were strongly suppressed by IL-10, showed enrichment of IRF1 motifs (Fig. 2g, Extended Data Fig. 2e). This suggests that genes that are regulated by IRFs may be more susceptible to inhibition by IL-10.

### IL-10 decreases chromatin accessibility and preferentially suppresses IRF activity

We next used ATACseq to test whether inhibition of H3K27ac by IL-10 was associated with decreases in chromatin accessibility, and footprinting under ATACseq peaks to identify changes in transcription factor occupancy. Surprisingly, IL-10 had a substantially more restricted effect on LPS-induced chromatin accessibility, suppressing only 1112 out of 8813 (12.6%) LPS-induced ATACseq peaks (Fig. 3a-d), compared to the broader suppression of 60% of LPS-induced H3K27ac peaks (Fig. 2a). Representative gene tracks of LPS-induced ATACseq peaks that were suppressed by, or resistant to, IL-10 are depicted in Fig. 3e. To gain insight into what may distinguish LPS-induced open chromatin regions that are closed versus remain open with IL-10 pretreatment we performed pathway enrichment analysis of genes associated with these peaks and motif enrichment analysis. Genes associated with suppressed peaks showed more significant enrichment in interferon pathways, whereas genes associated with both types of peaks were comparably enriched in NF-κB and inflammatory pathway genes (Extended Data Fig. 3a). LPS-induced ATAC peaks that were suppressed by IL-10 showed a similar suppression of H3K27ac (Extended Data Fig. 3b). Thus, a subset of genomic elements targeted by IL-10 show suppression of both activating histone marks and chromatin accessibility. Chromatin regions that lost accessibility were distinguished by enrichment of ISRE/IRF motifs (Fig. 3f), which are associated with canonical ISGs and bind IFN-activated transcription complex ISGF3 comprised of STAT1, STAT2 and IRF9 ^29^. Accordingly, IL-10-suppressed chromatin regions were associated with canonical ISGF3 target genes (Fig. 3e, right and Fig. 3g). These results indicate that IL-10 preferentially suppresses chromatin accessibility at ISG loci that are induced by ISGF3; however, ISREs can also bind IRFs such as IRF5 ^27, 39^,suggesting that suppression of IRFs can also contribute to loss of chromatin accessibility, including at NF-κB target genes (Extended Data Fig. 3a).

**Figure 3.**
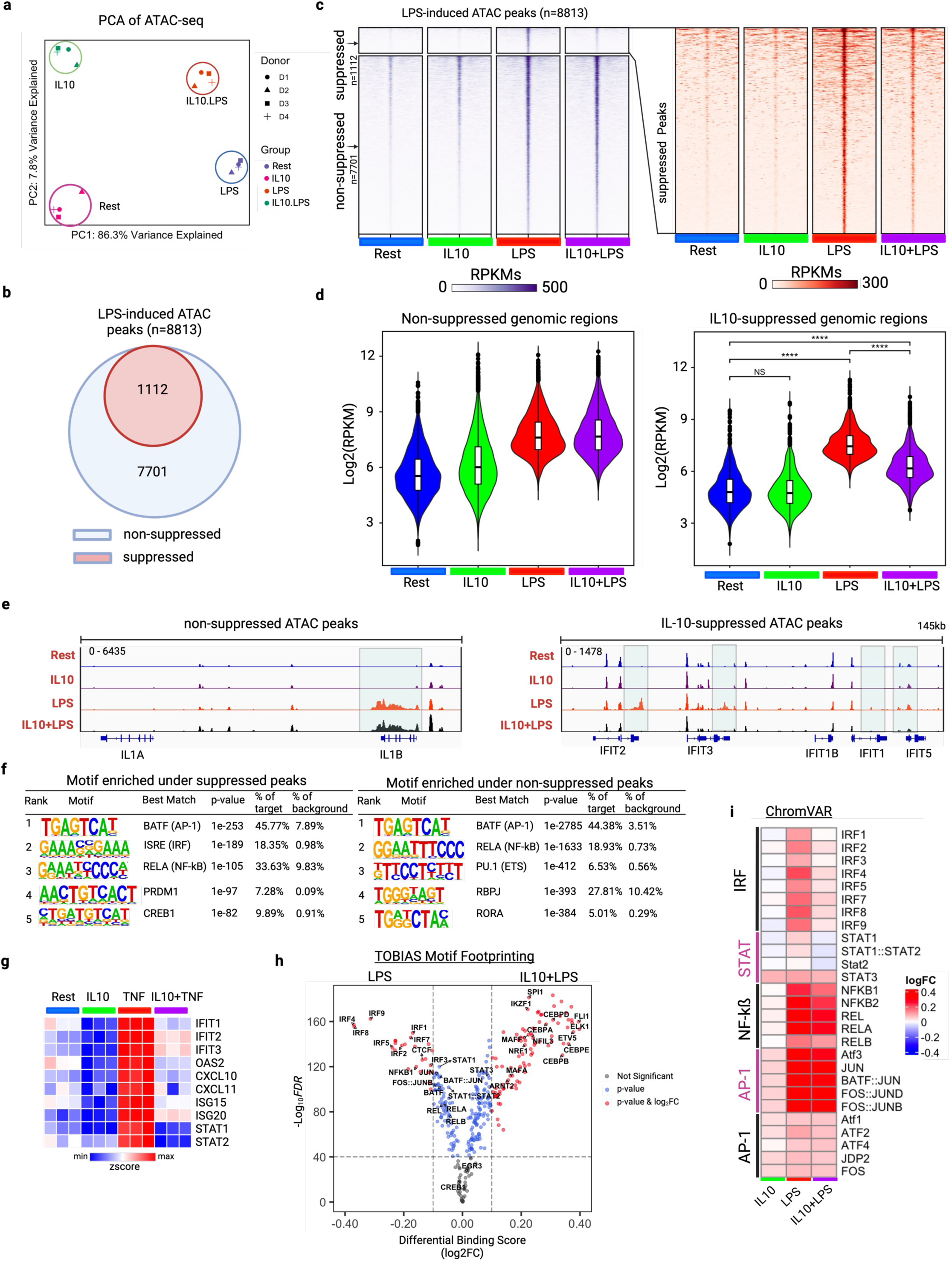
IL-10 decreases chromatin accessibility at a subset of enhancers and preferentially suppresses IRF factor activity. **a-i**, analysis of ATACseq data obtained using monocytes from 4 independent donors **a,** PCA plot of ATAC-seq data for Resting, IL-10, LPS, and IL-10+LPS treated primary human monocytes **b,** Venn diagram illustrating numbers of LPS-induced ATAC peaks that are either suppressed or not suppressed by IL-10 treatment. **c,** Heatmap of the normalized signal density of suppressed and non-suppressed LPS-induced ATAC peaks under indicated conditions. The results are presented in RPKM values within a range of ± 2.0kb around peak centers. **d,** Violin plots showing normalized average signal density of suppressed and non-suppressed LPS-induced ATAC peaks (corresponding to peaks in **c**) under indicated conditions. Data plotted as log2 (RPKM+1) counts of ATAC reads. ****p < 0.0001 by Two-way ANOVA with Tukey’s multiple comparisons test **e,** Representative IGV gene tracks of LPS-induced ATAC peaks that are non-suppressed (left panel, *IL1B* locus) and suppressed (right panel, *IFIT* locus) by IL-10. **f,** *De novo* motif analysis results using HOMER of LPS-induced ATAC peaks that are either suppressed or non-suppressed by IL-10. **g,** Heatmap depicting expression of LPS-induced ISGs associated with closing of chromatin by IL-10 treatment. **h,** Volcano plot of differential binding analysis of JASPAR motifs by TOBIAS using BINDetect algorithm for LPS versus IL-10+LPS treated human monocyte ATAC peaks. **i,** Heatmap of the differential TF activity scores derived from ChromVAR motif enrichment analysis of ATAC data for IL-10, LPS, and IL-10+LPS treated human monocytes, compared to resting control. The activity scores are illustrated for four major TF groups: IRFs, STAT, NF-κB, and AP-1.

We next performed footprinting analysis using TOBIAS, which measures occupancy of TFs motifs within ATACseq peaks and thus identifies which TFs are actually binding ^40^. As expected, LPS induced occupancy of NF-κB and AP-1 motifs, with a lesser occupancy of IRF motifs; IL-10 induced occupancy of STAT3 motifs (Extended Data Fig. 3c). In monocytes stimulated with IL-10 + LPS, occupancy of NF-κB and AP-1 motifs was mostly preserved, while occupancy of IRF motifs was diminished relative to LPS alone (Extended Data Fig. 3c, right panel). A direct comparison of the LPS versus IL-10 + LPS conditions confirmed that IL-10 most strongly and significantly suppressed IRF binding (Fig. 3h). We extended our motif footprinting analysis to TNF stimulation and uncovered a remarkable similarity in IL-10’s impact on TF motif occupancy, mirroring observations from LPS stimulation. Notably, TF occupancy of NF-κB and AP-1 motifs remained preserved, while occupancy of IRF motifs was distinctly suppressed (Extended Data Fig. 3d). Furthermore, a complementary computational analysis using ChromVAR that generates a TF activity score confirmed that IL-10 predominantly suppressed IRF activity, while LPS-induced NF-κB and AP-1 activity was mostly preserved in cells stimulated with IL-10 + LPS relative to LPS alone (Fig. 3i). The lack of suppression of LPS- and TNF-induced NF-κB and AP-1 activity by IL-10 is in accord with the literature showing that IL-10 does not suppress NF-κB or MAPK signaling ^11, 14^. Consistent with the HOMER motif enrichment analysis in Fig. 3f, ChromVAR also showed that IL-10 suppressed activity of STAT1-STAT2, which is induced by autocrine IFN-b. Overall, these results indicate that IL-10 suppresses the LPS-TLR4-induced transcriptional response mainly by targeting IRF family factors and suggest that IL-10 also interrupts the well-established TLR4-induced IFN-b-mediated autocrine loop ^29, 41^.

### IL-10 broadly suppresses LPS-induced IRF1 binding and associated histone acetylation

As IRF1 is induced by LPS and has been implicated in induction of both ISGs and inflammatory genes, we next investigated the effects of LPS and IL-10 on the genomic profile of IRF1 binding (Fig. 4a-b and Extended Data Fig. 4a-b). In accord with low level tonic IFN signaling in myeloid cells ^29, 42, 43^, unstimulated monocytes showed a small number of IRF1 binding peaks that were greatly induced by LPS. Strikingly, IL-10 strongly and significantly suppressed LPS-induced IRF1 binding broadly across the genome; 3931 out of 4395 LPS-induced peaks (89.5%) were lost with IL-10 pre-treatment (Fig. 4a and 4b). Pathway enrichment analysis of genes associated with LPS-induced IRF1 binding showed significant enrichment of IFN and inflammatory response pathways (Extended Data Fig. 4c). Motif analysis showed that LPS-induced IRF1-binding peaks were enriched for IRF motifs (as expected) and also AP-1 and NF-κB motifs (Fig. 4c). IRF1 binding peaks that were (partially) preserved under IL-10 stimulation conditions were enriched for IRF1 and AP-1 but not for NF-κB motifs, suggesting that genomic elements that bind both IRF1 and NF-κB are preferentially suppressed.

**Figure 4:**
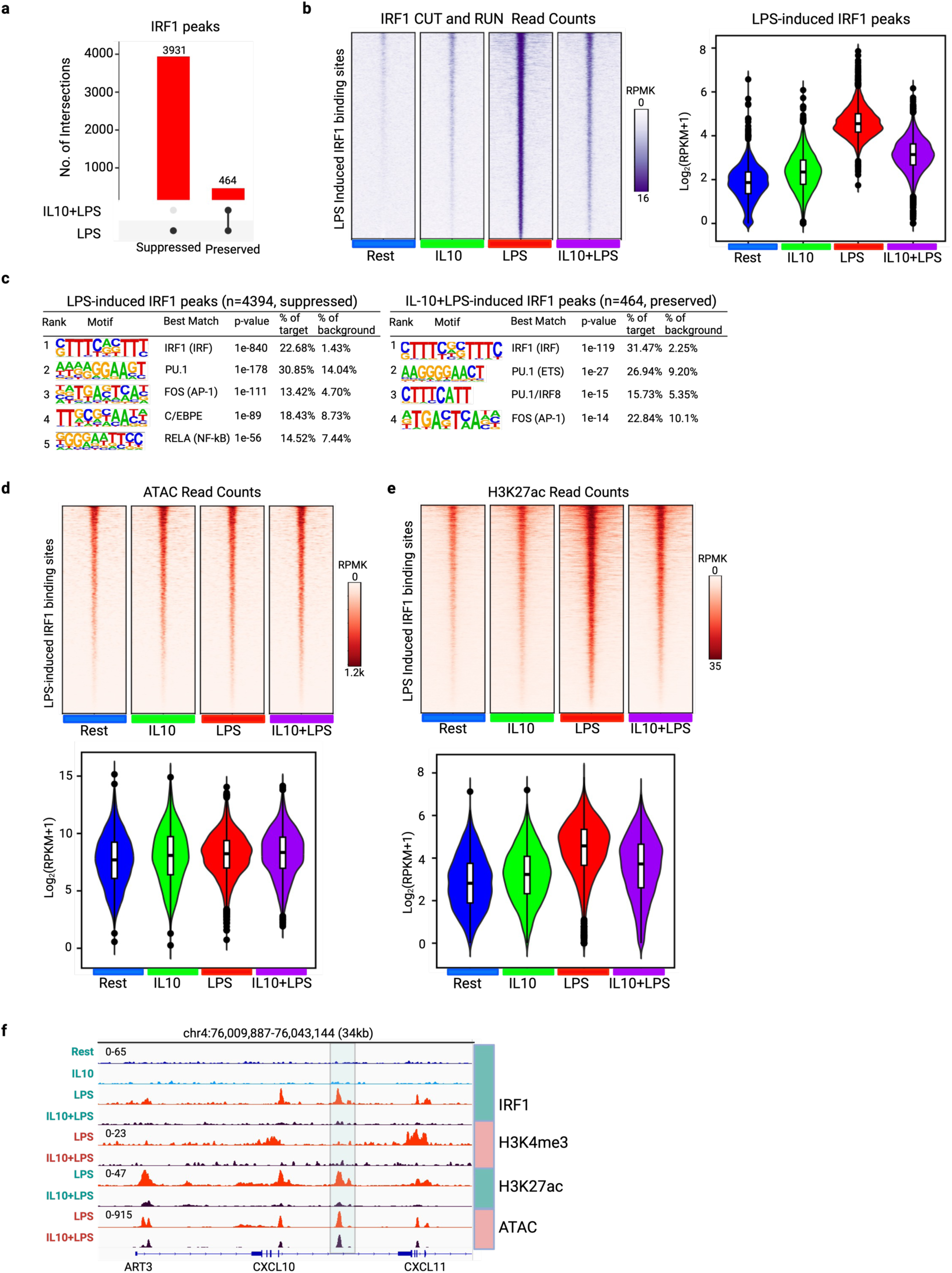
IL-10 broadly suppresses LPS-induced IRF1 binding and associated histone acetylation. **a-f,** analysis of IRF1 CUT and RUN data obtained using monocytes from 3 independent donors **a,** UpSet plot depiction of LPS-induced IRF1 peaks (Log2FC ≥ 1 and FDR ≤ 0.05) that were lost (unique) or preserved (common) after IL-10 treatment. **b,** Heatmap of the normalized signal density of LPS-induced IRF1 peaks (left panel) and violin plot showing normalized average signal density under indicated conditions (right panel) **c,** *De novo* motif analysis results using HOMER of IRF1 binding peaks, categorized as either unique to LPS (left panel) or common between LPS and IL-10+LPS (right panel). **d,** Heatmap of the ATAC normalized signal density surrounding LPS-induced IRF1 peaks (top panel), and violin plot showing normalized average signal density under indicated conditions (bottom panel). **e,** Heatmap of the H3K27ac normalized signal density surrounding LPS-induced IRF1 peaks (top panel), and violin plot showing normalized average signal density under indicated conditions (bottom panel). **f,** Representative IGV gene tracks illustrating LPS-induced IRF1 binding, H3K4me3 and H3K27ac peaks, and ATAC peaks at the *CXCL10* locus. **b, d and e**, heatmap were generated using RPKM values within in a range of ± 2.0kb around peak centers and for violin plot data are plotted as log_2_(RPKM+1) counts.

We next investigated whether LPS-induced IRF1 binding was associated with increases in chromatin accessibility or chromatin activation, as assessed by increased H3K27ac. Interestingly, chromatin regions bound by IRF1 were already accessible in resting monocytes, and accessibility minimally changed after IL-10 or LPS stimulation (Fig. 4d). This observation aligns with our previous finding that IL-10 pretreatment strongly suppressed IRF footprints with lesser effects on chromatin accessibility (Fig. 3). In contrast, IRF1 binding peaks were characterized by LPS-inducible H3K27ac, which was strongly decreased when IL-10 suppressed IRF1 binding (Fig. 4e). Examples of IL-10-mediated suppression of IRF1 binding, and of H3K4me3 and H3K27ac, while open chromatin is preserved are depicted in representative gene tracks in Fig. 4f (boxed peak) and Extended Data Fig. 4d). These results suggest that IRF1 does not regulate chromatin accessibility, but instead is associated with activation of chromatin via histone acetylation, and the latter is suppressed by IL-10.

### IL-10 alone induces limited suppressive epigenomic reprogramming

One potential mechanism of IL-10 action is reprogramming of the monocyte epigenome such that a subsequent LPS-induced signal, even if intact, encounters a closed and suppressive chromatin environment that prevents transcription activation. Thus, we analyzed the effects of IL-10 alone on gene expression, chromatin states and IRF1 binding. RNAseq analysis showed induction of well-known IL-10 target genes (Fig. 5a). Pathway analysis of IL-10-supressed genes revealed most significant suppression of IFN response genes (Fig. 5b), which is consistent with suppression of tonic IFN signaling that maintains basal expression of STAT1 and select ISGs ^42^. Accordingly, IL-10 suppressed basal *STAT1* expression; however, basal *IRF1* expression was not suppressed (Figs. 5a and 6a). IL-10 had minimal effects upon low basal expression of inflammatory genes and minimally suppressed basal H3K27ac (Fig. 5c) or basal IRF1 binding (Fig. 4b), and suppressed basal chromatin accessibility at only 477 out of 4811 ATACseq peaks (Fig. 5d). *De novo* motif enrichment analysis under the IL-10-suppressed ATACseq peaks showed enrichment of IRF motifs (Fig. 5e) and TOBIAS footprinting analysis showed decreased occupancy of STAT1 and IRF9 sites (Extended Data Fig. 3c). In contrast to suppressed peaks, IL-10-induced ATACseq and H3K27ac peaks were enriched for AP-1 and STAT3 motifs (as expected) (Fig. 5e and 5f). Collectively, these results suggest that IL-10 interrupts tonic IFN signaling via ISGF3 that maintains expression of STAT1 and select ISGs. However, the different regulation of IRF1 binding and H3K27ac, along with suppression of completely distinct ATACseq peaks by IL-10 under basal versus LPS-stimulated conditions (Fig. 5g) indicate that IL-10 utilizes additional mechanisms to suppress the LPS response. Furthermore, limited suppression of accessible chromatin regions and lack of suppression of H3K27ac suggest that IL-10 does not broadly remodel the epigenome in resting monocytes to make genes resistant to induction by LPS.

**Figure 5:**
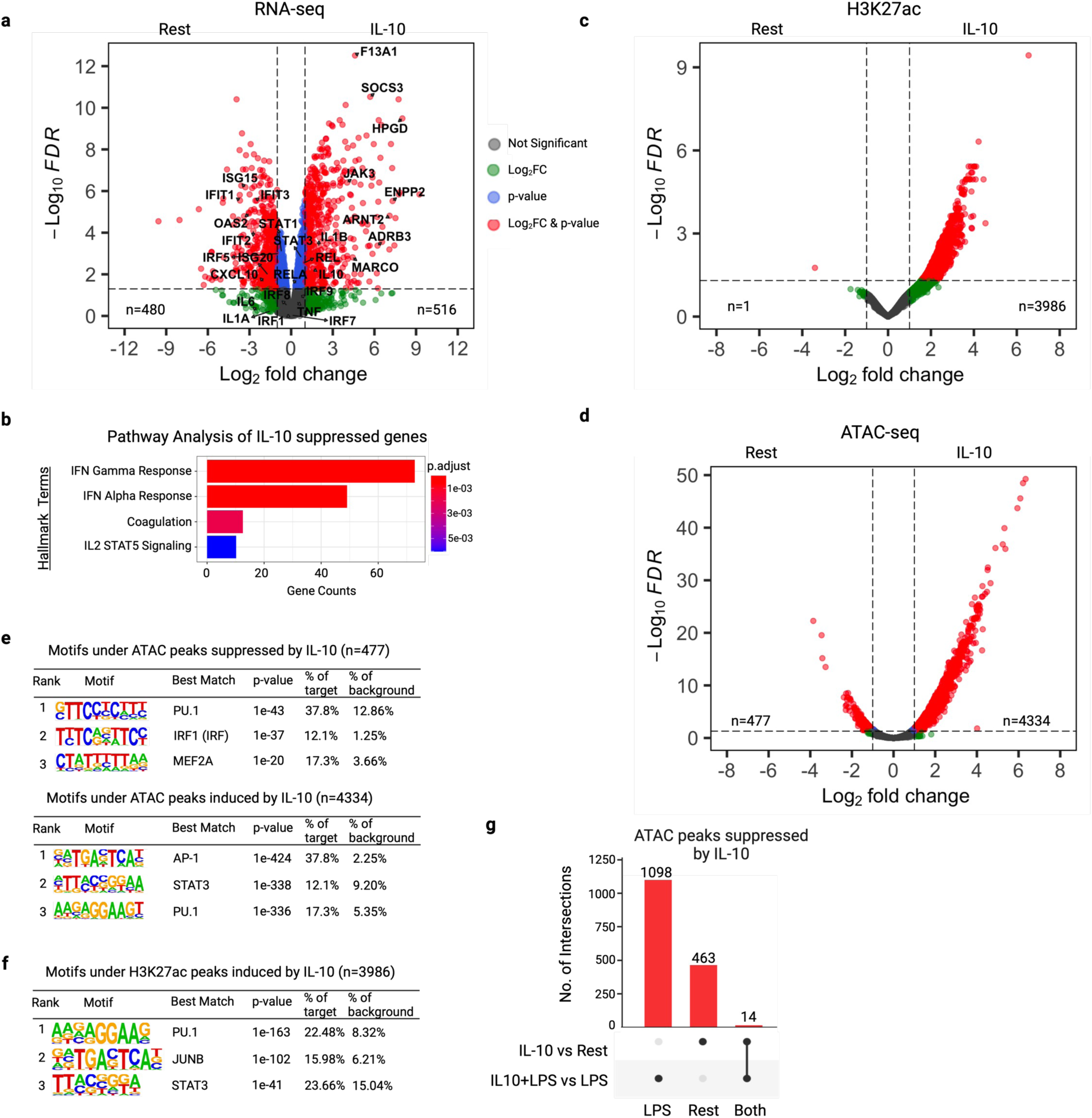
IL-10 alone induces limited epigenomic reprogramming. **a,** RNAseq. Volcano plot of DEGs regulated by IL-10 (100 ng/mL, up=516 and down=480) after 18h treatment of human primary monocytes relative to resting control (n=3 independent donors). **b,** Hallmark pathway enrichment analysis of IL-10 suppressed genes. **c,** Volcano plot of H3K27ac peaks that are regulated by IL-10 (up=3986, down=1) treatment of human primary monocytes (n= 2 independent donors). **d,** Volcano plot of ATAC peaks that are regulated by IL-10 (up=4334, down=477) treatment of human primary monocytes (n= 4 independent donors). **e-f,** *De novo* motif analysis results using HOMER **e,** on ATAC peaks categorized as IL-10 suppressed peaks (top panel) and IL-10-induced peaks (bottom panel), **f,** on H327ac binding peaks induced by IL-10. **g,** UpSet plot showing minimal overlap between ATAC peaks suppressed by IL-10 in resting monocytes and in monocytes stimulated with LPS.

**Figure 6:**
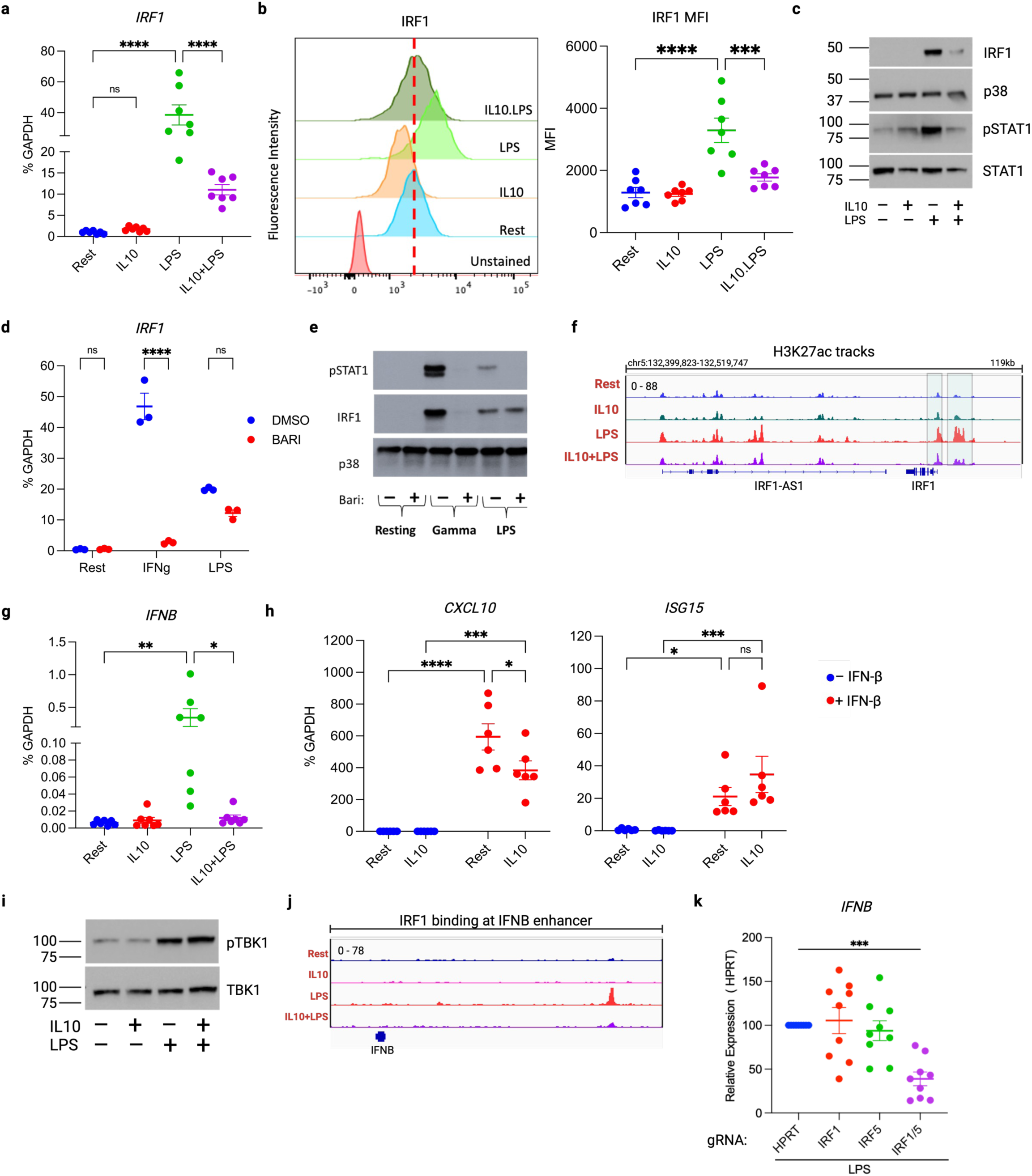
IL-10 suppresses IRF1 and IRF5 and the TLR4-induced IFN-b-mediated autocrine loop. **a,** IRF1 mRNA was measured by qPCR and normalized relative to GAPDH mRNA in cells stimulated as indicated (n=7 independent donors). Data are depicted as mean ± SEM. ****p < 0.0001 by One-way ANOVA with Tukey’s multiple comparisons test. **b,** FACS to analyze protein level of IRF1 in human monocytes in the indicated conditions. Left panels, representative histogram plot; right panels, MFI of FACS experiments (n=7 independent donors). **c,** Immunoblot of IRF1, p38, pSTAT1 and STAT1 using whole cell lysates of cells treated with indicated stimuli (n = 3 independent donors, p38 used as loading control). **d,** IRF1 mRNA was measured by qPCR and normalized relative to GAPDH mRNA in cells pre-treated with baricitinib for 30min prior to stimulation with IFNγ or LPS for 3hours (n=3 independent donors, IFNγ used as a control). **e,** Immunoblot of IRF1, pSTAT1 and p38 using whole cell lysates of cells treated with baricitinib for 30min prior to stimulating with IFNγ or LPS for 3hours (representative blot from one out of 3 independent donors, p38 used as loading control) **f,** IGV gene track of LPS-induced H3K27ac peak at IRF1 locus. **g,** mRNA of *IFNB* was measured by qPCR and normalized relative to GAPDH mRNA in cells stimulated with indicated stimuli (n=7 independent donors). **h,** mRNA of indicated genes was measured by qPCR and normalized relative to GAPDH mRNA in cells treated with IL-10 for 18 hours and stimulated with exogenous IFN-b to assess whether IL-10 blocks IFN-b signaling (n=6 independent donors). **i,** Immunoblot of phospho-TBK1 and TBK1 using whole cell lysates of cells treated with indicated stimuli (representative blot from one out of 3 independent donors). **j,** IGV gene tracks of LPS-induced IRF1 binding at *IFNB* enhancer. **k,** mRNA of *IFNB* was measured by qPCR and normalized relative to GAPDH mRNA after CRISPR-Cas9-mediated editing of IRF1, IRF5 or IRF1+IRF5 together (IRF1/5) and stimulation with LPS for 3h (n=9 independent donors). for **a, b, d, g, h, and k**, each dot represents an independent donor and data are depicted as mean ± SEM. *p < 0.05; **p < 0.001; ***p ≤ 0.0005; ****p < 0.0001 by One-way ANOVA with Tukey’s multiple comparisons test (**a, b, d, g, and k**) and two-way ANOVA with Tukey’s multiple comparison test (**h**).

### IL-10 suppresses IRF1 and IRF5 to inhibit TLR4-induced *IFNB* expression

We wished to understand how IL-10 suppresses IRF1, and the functional consequences for induction of the LPS-induced IFN-b-mediated autocrine loop. IL-10 strongly suppressed LPS-mediated induction of IRF1 at both mRNA and protein levels (Fig. 6a-c). IRF1 is best known as an ISG; however, the Jak inhibitor baricitinib, which strongly suppressed induction of IRF1 by IFNγ, only weakly affected induction of IRF1 by LPS (Fig. 6 d and 6e). Thus, under our experimental conditions, IRF1 was primarily induced by LPS via non-IFN-mediated pathways and was associated with decreased H2K27ac at an enhancer in the *IRF1* locus (Fig. 6f). These results show that IL-10 inhibits IRF1 mainly by suppressing its induction, which is only partially mediated by autocrine IFN-b signaling.

Induction of ISGs by TLR4 or TNFR signaling requires production of IFN-b, which binds to its receptor IFNAR to activate Jak-STAT signaling via tyrosine phosphorylation of STAT1 and STAT2 and their association with IRF9 in the ISGF3 complex. The inhibition of STAT1 activation and ISG induction by IL-10, as observed in Fig. 6c and Fig. 1, respectively, could occur via suppression of IFN-b production, suppression of signaling by IFN-b, direct suppression of ISG transcription, or a combination thereof. IL-10 significantly suppressed LPS-induced *IFNB* expression (Fig. 6g and Extended Data Fig. 5a) but had minimal effects on ISG induction by exogenous IFN-b (Fig. 6h and Extended Data Fig. 5b), suggesting a role for inhibition of IFN-b production in the decreased expression of ISGs. As induction of IFN-b by TNF is dependent on IRF1 ^36, 37^, suppression of TNF-induced IRF1 by IL-10 (Fig. Extended Data Fig. 5c) explains at least in part how IL-10 inhibits induction of ISGs by TNF. In contrast, IFN-b induction by TLR4 is generally thought to mediated by TBK-IRF3 signaling, which has been established in mouse myeloid cells ^44, 45, 46^. However, in human monocytes IL-10 did not affect TBK1 activation (Figs. 6i and Extended Data Fig. 5d) and we considered the possibility that IRF1 mediates *IFNB* induction in human monocytes. This notion was supported by strong LPS-induced IRF1 binding at an enhancer at the *IFNB* locus which was suppressed by IL-10 (Fig. 6j). To directly test the role of IRF1 in regulating *IFNB* gene expression, we utilized CRISPR-Cas9 to edit the *IRF1* gene locus in primary human monocytes, which resulted in an almost complete loss of IRF1 protein expression (Extended Data Fig. 5d). As experiments with multiple blood donors using monocytes with disrupted *IRF1* did not show a significant decrease in LPS-induced *IFNB* induction (Fig. 6k), we addressed the possibility of redundant IRF function. Strikingly, combined CRISPR-mediated disruption of IRF1 and IRF5, which was also suppressed by IL-10 (Extended Data Fig. 5e) highly significantly decreased LPS-induced *IFNB* mRNA (Fig. 6k; p < 0.0005, mean 61.07% reduction). These results show important roles for IRF1 and IRF5 in *IFNB* induction by LPS and support the idea that IL-10 suppresses *IFNB* induction in human monocytes at least in part by targeting IRF1 and IRF5. A role for IRF1 in *IFNB* induction in human monocytes is in line with results in the initial report of the molecular cloning of IRF1 ^32^, the role of IRF1 in TNF-induced *IFNB* expression ^36, 37^, and a more recent report proposing a role for IRF1 in enabling IRF3 function ^47^.

### Inflammatory and interferon-stimulated genes suppressed by IL-10 are IRF1/5 targets

We wished to more broadly characterize the functional consequences of disruption of *IRF1* and *IRF5* for the LPS-induced gene response in human monocytes. In line with decreased *IFNB* expression (Fig. 6k), editing of *IRF1* and *IRF5* significantly decreased LPS induction of ISGs *CXCL10* and *ISG15* (Fig. 7a). Interestingly, editing of *IRF1* and *IRF5* also significantly decreased LPS-induced expression of inflammatory genes *TNF* and *IL6* (Fig. 7b). These results provide strong genetic evidence for a key role of IRF1 and IRF5 in LPS-induced gene expression, which complements the epigenomic data presented above for IRF1.

**Figure 7:**
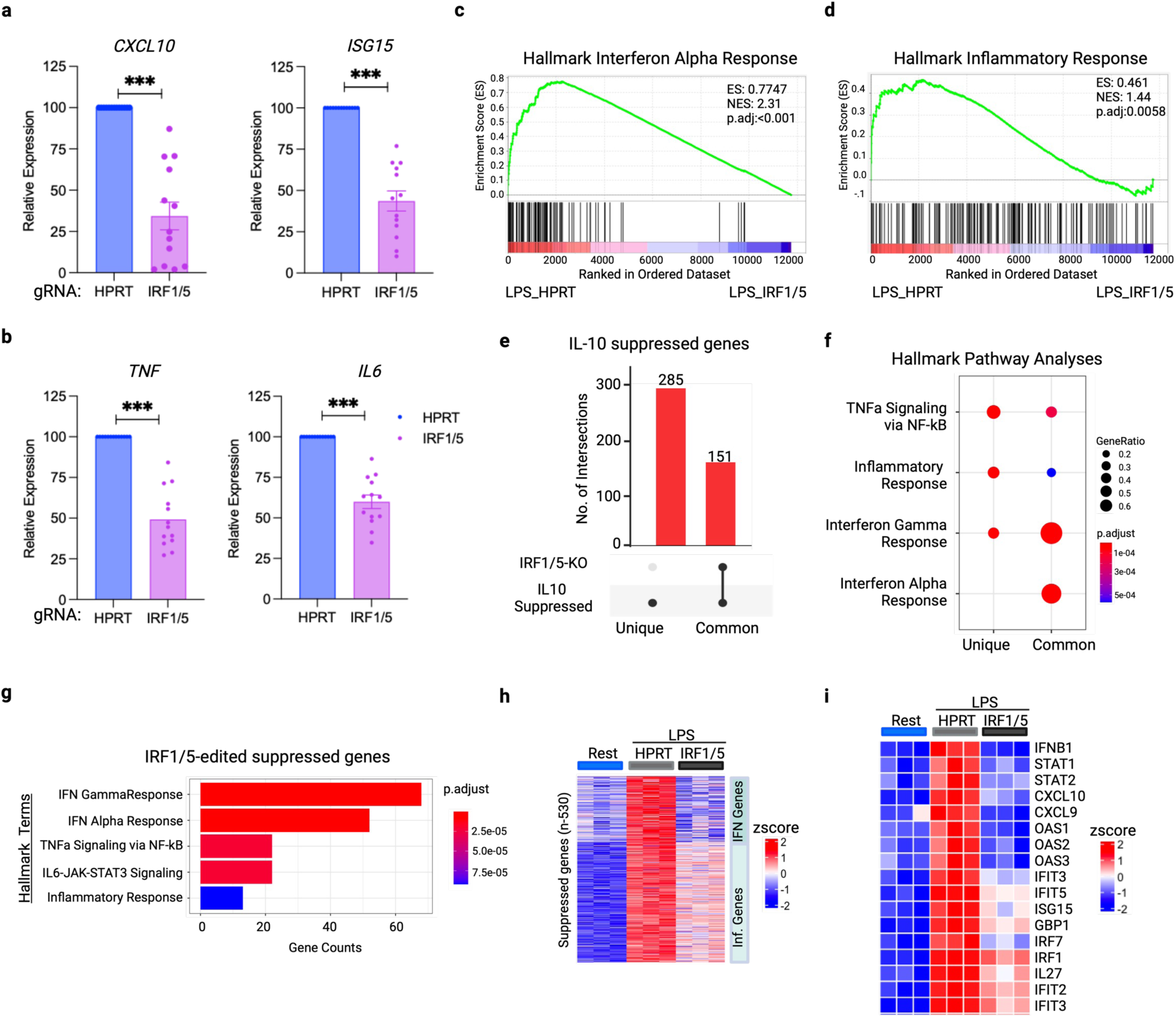
IRF1/5 target genes are suppressed by IL-10 in human monocytes. **a-b,** mRNA of indicated **a,** IFN-response genes (*CXCL10* and *ISG15*) and **b,** inflammatory genes (*TNF* and *IL-6*) was measured by qPCR and normalized relative to GAPDH mRNA. This analysis was conducted after CRISPR-Cas9 gRNA targeting of *IRF1*+*IRF5* or *HPRT*-control for 48 hours, followed by stimulation with LPS for 3 hours (n=13 independent donors). Data are depicted as mean ± SEM. ***p < 0.001 by Wilcoxon matched-pairs signed rank test. **c-d,** GSEA plots of RNA-seq data performed on DEGs ranked by log2CPMs in *IRF1*+*IRF5*-edited vs. control *HPRT*-edited primary human monocytes stimulated with LPS for 3 hours (n=3 independent donors). **c,** Hallmark Interferon Alpha Response enrichment plot. **d,** Hallmark Inflammatory Response enrichment plot. **e,** UpSet plot showing overlap between LPS-induced genes suppressed by IL-10 (as shown in Fig. 1) and genes dependent upon IRF1/5 as identified by RNAseq as in **c, d**. **f-g,** Hallmark pathway enrichment analysis **f,** of genes associated with groups identified in Fig. 7e. and **g,** of genes downregulated in IRF1/5-edited primary human monocytes compared HPRT-edited. **h-i,** Heatmaps of **h,** genes selectively downregulated in IRF1/5-edited monocytes compared to control HPRT-edited monocytes, clustered by Inflammatory and IFN response genes **i,** representative IRF1/5-dependent interferon and inflammatory response genes.

We reasoned that if IL-10 suppressed gene induction in part by suppressing IRF1 and IRF5 expression, then the effects of IRF1/IRF5 disruption should at least partially phenocopy the effects of IL-10 on LPS-induced gene expression. This notion was tested by using RNAseq to identify LPS-induced genes that were dependent on IRF1/5 (Extended Data Fig. 6a) and comparing this gene set to the IL-10 suppressed gene set defined above in Figure 1. Similar to analysis of IL-10 suppressed genes, GSEA comparing gene expression between the LPS-stimulated control or IRF1/5-edited monocytes showed most significant suppression of IFN response genes, and highly significant suppression of Hallmark inflammatory genes (Fig. 7c-d). We then use a similar strategy as described in Fig. 1 and Extended Data Fig. 1 for IL-10-suppressed genes to identify a core set of 530 LPS-induced genes that were dependent on IRF1/5 (Extended Data Fig. 6b-f). Interestingly, 151 out of 436 IL-10-suppressed genes were dependent on IRF1/5 (Fig. 7e). This suggests that approximately 35% of the gene suppressive activity of IL-10 is mediated by targeting IRF1/5, and, importantly, the IL-10-suppressed genes that are IRF1/5 targets are highly significantly enriched in both IFN response and inflammatory genes (Fig. 7f-g). Similar to IL-10-suppressed genes, IRF1/5-dependent genes segregated into genes whose expression was strongly or partially diminished when IRF1/5 were disrupted, with ISGs being more strongly suppressed (Fig. 7h-i). These results identify IRF1/5 targets in human monocytes and implicate targeting of IRF1/5 as a key mechanism by which IL-10 suppresses LPS-induced IFN and inflammatory responses.

## Discussion

Although the importance of IL-10 in restraining inflammation and suppressing inflammatory diseases is established, mechanisms by which IL-10 inhibits myeloid cell activation have remained elusive, despite decades of research. In this study, we applied an integrated epigenomic analysis to define mechanisms by which IL-10 inhibits activation of monocytes by the prototypical potent TLR ligand LPS. We found that instead of inhibiting core TLR4-activated pathways such as NF-κB and MAPK signaling, IL-10 targets expression and DNA binding/transcription factor activity of IRF proteins that are activated or induced by TLR4. This resulted in highly significant inhibition of inflammatory NF-κB target genes, in whose activation IRFs play an amplifying role, and a near-complete suppression of ISGs that are IRF-dependent. Mechanisms of TLR4 target gene inhibition included downregulation of chromatin accessibility and de novo enhancer formation at ISGs, and decreased induction of IRF1-associated H3K27ac activating histone marks at both ISGs and inflammatory gene loci. IL-10 suppressed TNF-induced ISG and inflammatory gene expression in a similar manner, further supporting an epigenetic mechanism of action. These results provide a mechanism by which IL-10 suppresses induction of inflammatory genes describe an underappreciated suppression of IFN responses by epigenetic mechanisms.

The IRF family of transcription factors has been most closely linked to IFN responses ^29, 48, 49, 50^. IRF1, IRF8 and IRF9 were initially described as interferon-inducible genes and they cooperate with IFN-activated STATs to amplify and extend induction of ISGs, many of which contain binding sites for both STATs and IRFs. In contrast, IRF3 and IRF7 are activated by TLRs and nucleic acid-sensors via phosphorylation and activate expression of *IFNB* and *IFNA* genes to drive downstream IFN responses. IRF5 is activated by phosphorylation similarly to IRF3/7 but predominantly targets inflammatory genes, likely in cooperation with NF-κB ^27, 51, 52^. The current model for IRF protein functions in the TLR4 response is based mostly on research with mouse myeloid cells such as in vitro differentiated bone marrow-derived macrophages, and posits that activation of IRF3 via the signaling adaptor protein TRIF plays a key role in inducing *IFNB*, which in turn activates ISGF3 (comprised of STAT1/STAT2/IRF9) to induce ISGs ^46, 53, 54^. These ISGs include IRF1 that cooperates with ISGF3, and IRF9 that increases ISGF3 amounts, to further drive ISG expression and the IFN response. In contrast, signaling via adaptor MyD88 activates NF-κB and MAPK-AP1 to induce inflammatory genes, and also IRF5 which amplifies inflammatory gene induction and can contribute to induction of ISGs. In contrast to this model, our results in primary human monocytes highlight a prominent role for TLR4-induced IRF1, in cooperation with IRF5, in direct induction of *IFNB*, ISGs and inflammatory genes (Extended Data Fig. 7). Combined disruption of *IRF1* and *IRF5* did not completely abrogate gene induction, and it is possible that additional IRFs contribute to gene induction. As IRF1 expression is highly regulated, these results suggest that IRF1 performs a rheostat function in broadly determining the magnitude of TLR4-induced gene responses, and this rheostat is tuned by modulating IRF1 expression (Extended Data Fig. 7).

The most novel and important finding of our study is that a key mechanism of IL-10 action is to turn down the IRF1-mediated ‘rheostat’, thereby turning off the IRF1-mediated amplification of TLR4- and TNF-induced IFN and inflammatory responses. IL-10 likely inhibits induction of IRF1 in part by disrupting an auto-amplification loop (Extended Data Fig. 7), which is a common mechanism for transcription factor induction ^55, 56^. Suppression of other IRF family members, in particular IRF5, contributes to IL-10 bioactivity. The consequences of this suppression of IRFs include loss of the activating H3K27ac mark at ISGs and inflammatory genes, loss of chromatin accessibility predominantly at ISGs, and loss of *IFNB* induction and activation of ISGF3. ISGF3 binding has been closely linked with chromatin opening and remodeling ^50^, whereas IRFs are associated with recruitment of histone acetyl-transferases (HATs) and histone acetylation ^48, 49^. Thus, the decrease of LPS-induced chromatin accessibility and H3K27ac at ISG loci is most likely explained by combined loss of IRF1, ISGF3, and likely additional IRF binding, and results in near-complete loss of gene induction. The decrease in H3K27ac at inflammatory gene loci is associated with decreased binding of IRF1 and likely other IRFs, but chromatin accessibility was preserved, and gene induction was decreased to a lesser extent than induction of ISGs. Preserved chromatin accessibility at these loci suggests that in our system IRFs bind to already open chromatin, and their function is related more to posttranslational modification of histones.

Two unexpected findings of our study are the important role of IRF1, together with IRF5, in induction of *IFNB* by TLR4, and the stronger suppression of ISGs than inflammatory genes by IL-10 in monocytes. We believe these two findings are related, as suppression of IRF-mediated *IFNB* induction will have a major effect on ISGs, while IRFs play more of an auxiliary role in inflammatory gene induction. Although the current paradigm is that IFN-b induction by TLR4 is essentially entirely dependent on IRF3, previous reports provided signaling and genetic evidence that human *IFNB* and mouse *Ifnb* gene induction by TNF is mediated by IRF1, and IRF1 deficiency abrogates TNF-induced IFN responses in vivo ^36, 37^. Thus, IRF1 has a physiological and context-dependent role in IFN-b induction. Furthermore, early studies reporting the molecular cloning of IRF1 suggested a role in IFN-b induction ^34^, and investigation of *IFNB* gene regulation implicated IRF1 interactions with NF-κB and AP-1/ATF factors in cooperative assembly of an enhanceosome complex and recruitment of chromatin-modifying enzymes ^57, 58^. As TBK1-IRF3 signaling remained intact in IL-10-treated monocytes, it is likely that IRF3 contributes to the residual induction of ISGs when IRF1 and IRF5 are abolished. It also remains possible that suppression of additional IRFs by IL-10 can contribute to its effects on LPS-induced gene activation.

Differences in the utilization of IRFs and ‘wiring’ of signaling pathways leading from TLR4 to inflammatory and IFN response genes can explain cell-type and context-dependent differences in the effects of IL-10 on TLR4-induced gene expression. The predominant utilization of IRF3 for TLR4-mediated *Ifnb* induction in mouse macrophages may explain why a role for IRF1 was not previously fully appreciated. Conversely, an important role for IRF1 in human monocytes, where it binds to *IFNB*, ISGs and inflammatory genes and is suppressed by IL-10, contributes to the suppression of both ISGs and inflammatory genes by IL-10 in these cells. An attractive potential explanation for cell type-specific differences in IRF1-mediated *IFNB* induction that merits future investigation is differences in formation and function of an upstream IRF1 binding enhancer at the *IFNB* locus, whereas IRF3 may exert its effects primarily at the gene promoter. As IRF1 gene expression is rapidly and highly induced by IFNs, cytokines such as TNF, and inflammatory factors, these results suggest that monocytes are wired to rapidly amplify inflammatory and IFN responses. Similarly, monocytes activate the NLRP3 inflammasome by distinct mechanisms from macrophages, most notably bypassing the need for a “second signal” to release large amounts of mature IL-1b ^59, 60, 61^. This ability to ramp up inflammatory and IFN responses is consonant with the physiological function of monocytes, whose activation upon entering infected tissues is of paramount importance for host defense ^62, 63^. Although it is possible that differences in the IRF1 amplification loop are related to species differences between human and mouse cells, we believe that these differences more likely reflect changes that occur during cell differentiation, and thus have not been previously observed as the small numbers of mouse blood monocytes has precluded these studies. The view that TLR signaling and utilization of IRFs can change during myeloid cell differentiation is supported by signaling differences between distinct mouse myeloid cell subsets, and by changes in IRF activity and utilization that occur during in vitro differentiation of human monocytes to macrophages ^47, 64^.

A key role for IRF1 in host defense against various intracellular pathogens is well established in mouse models and human infections ^65, 66, 67, 68^. There is accumulating evidence supporting increased IRF1 expression and function in autoimmune diseases such as rheumatoid arthritis (RA) and systemic lupus erythematosus (SLE). Activated macrophages in inflamed RA joints show elevated IRF1 expression and increased expression of genes associated with IRF1-binding enhancers ^69^. IRF1 expression is also elevated in SLE monocytes, and its genome-wide binding profile is associated with histone acetylation and increased gene transcription ^70, 71, 72, 73^. Interestingly, both RA and SLE are characterized by a relative deficit in IL-10 expression or activity ^74, 75, 76, 77, 78^ suggesting that decreased inhibition of the IRF1 amplification loop may contribute to pathogenesis. Interestingly, IL-10 expression is suppressed by IRF5 and possibly by IRF1 ^79, 80^ suggesting cross regulation between IRFs and IL-10 and the importance of balanced activity of an IRF-IL-10 axis for proper regulation of inflammation and return to homeostasis. Together with our work, these studies suggest investigation of the IRF-IL-10 axis in autoimmune diseases as a fruitful area for future research.

In summary, this study explores the role of IL-10 in modulating inflammation and immune responses by focusing on its impact on gene expression in response to TLR/TNFR ligands in human monocytes. Utilizing epigenomic analysis techniques, we show that IL-10 targets IRF5 and an IRF1-mediated amplification loop, which are crucial for the induction of IFNB, ISGs, and other inflammatory genes. Our paper highlights the dualistic nature of immune regulation, with IRF1/5 driving the immune response to threats and IL-10 serving as a critical brake on this response. This balance between activation and inhibition underscores the complexity of immune regulation and points to potential therapeutic targets for controlling inflammatory and autoimmune diseases. The findings emphasize the importance of IRF1/5 as key players in the immune response and illustrate how its activity is modulated by IL-10 to suppress inflammatory gene induction.

## METHOD DETAILS

### Primary human monocytes

Deidentified buffy coats were purchased from the New York Blood Center following a protocol approved by the Hospital for Special Surgery Institutional Review Board. Peripheral blood mononuclear cells (PBMCs) were isolated using density gradient centrifugation with Lymphoprep (Accurate Chemical) and monocytes were purified with anti-CD14 magnetic beads from PBMCs immediately after isolation as recommended by the manufacturer (Miltenyi Biotec). Monocytes were cultured overnight at 37°C, 5% CO_2_ in RPMI-1640 medium (Invitrogen) supplemented with 10% heat-inactivated defined FBS (HyClone Fisher), penicillin-streptomycin (Invitrogen), L-glutamine (Invitrogen) and 20 ng/ml human M-CSF. Then, the cells were treated as described in the figure legends.

### Analysis of mRNA amounts (qPCR)

Total RNA was isolated using the RNeasy Mini Kit (QIAGEN, Cat#: 74106 ) following the manufacturer’s instructions. Reverse transcription of RNA into complementary DNA (cDNA) was performed using the RevertAid RT Reverse Transcription Kit (Thermo Fisher Scientific, Cat# K1691:) according to the manufacturer’s protocol, and the resulting cDNA was used for downstream analysis. For quantitative real-time PCR (qPCR), Fast SYBR Green Master Mix (Applied Biosystems, Cat#: 4385618) and a QuantStudio5 Real-time PCR system (Applied Biosystems) were used. CT values obtained from qPCR were normalized to the housekeeping gene *GAPDH*. Relative expression of target genes were calculated using the ΔCt method, where ΔCt represents the difference in threshold cycle values between the target gene and *GAPDH*. The results are presented as a percentage of *GAPDH* expression (100/2^ΔCt). Primer sequences used for the quantitative RT-qPCR reactions are provided in the Supplementary Table 1.

### Western blotting

For protein analysis we followed a previously reported method ^61^. Briefly, 2 x10^6^ human monocytes were washed with cold PBS after indicated treatments and harvested in 50uL cold lysis buffer containing Tris-HCl pH 7.4, NaCl, EDTA, Triton X-100, Na3VO4, phosSTOP EASYPACK, Pefabloc, and EDTA-free complete protease inhibitor cocktail. After a 10-minute ice incubation, cell debris was pelleted at 16,000xg at 4°C for 10 minutes. The soluble protein fraction was combined with 4× Laemmli Sample buffer (Bio-RAD, Cat#: 1610747) containing 2-mercaptoethanol and subjected to SDS-PAGE (electrophoresis) on 4–12% Bis-Tris gels. Following transfer of gels to polyvinylidene difluoride membranes, membranes were blocked in 5% (w/v) Bovine Serum Albumin in TBS with Tween-20 (TBST) at room temperature for at least one hour. Incubation with primary antibodies (diluted 1:1000 in blocking buffer) occurred overnight at 4°C. Membranes were washed three times with TBST and probed with anti-rabbit IgG secondary antibodies conjugated to horseradish peroxidase (diluted 1:2000 in blocking buffer) for one hour at room temperature. Enhanced chemiluminescent substrates (ECL western blotting reagents (PerkinElmer, cat: NEL105001EA) or SuperSignal West Femto Maximum Sensitivity Substrate (Thermo Fisher Scientific, Cat: 34095) were used for detection, followed by visualization on autoradiography film (Thomas Scientific, cat: E3018). For multi-protein detection on the same experimental filter while minimizing stripping and reprobing, membranes were horizontally cut based on molecular mass markers and the target proteins’ sizes. Restore PLUS western blotting stripping buffer (Thermo Fisher Scientific, Cat#: 46430) was applied for membranes requiring multiple primary antibody probes. Antibodies are listed in Supplementary Table 2.

### RNA sequencing

Libraries for sequencing were prepared using mRNA that was enriched from total RNA using NEBNext® Poly(A) mRNA Magnetic Isolation Module (New England Biolabs (NEB), Cat#: E7490L), and enriched mRNA was used as an input for the NEBNext Ultra II RNA Library Prep Kit (NEB, Cat#: E7770L), following the manufacturer’s instructions. Quality of all RNA and library preparations was evaluated with BioAnalyser 2100 (Agilent). Libraries were sequenced by the Genomic Resources Core Facility at Weill Cornell Medicine using a Novaseq SP flow cell, 50-bp pair-end reads to a depth of ∼20 - 40 million reads per sample. Read quality was assessed and adapters trimmed using FastQC and cutadapt. Reads were then mapped to the human genome (hg38) and reads in exons were counted against Gencode v38 with STAR Aligner. Differential gene expression analysis was performed in R using edgeR. Only genes with expression levels exceeding 4 counts per million reads in at least one group were used for downstream analysis. Benjamini-Hochberg false discovery rate (FDR) procedure was used to correct for multiple testing. Genes were categorized as upregulated if log2FC ≥ 1 and FDR ≤ 0.05 threshold was satisfied, downregulated if log2FC ≤ -1 and FDR ≤ 0.05. Heatmap with K-mean clustering was done using Morpheus web application and replotted using the R package pheatmap. Bioinformatic tools used are listed in Supplementary Table 3.

### Gene Set Enrichment Analysis (GSEA)

GSEA was conducted using the Broad Institute’s GSEA software (version 4.3.2). Log2 transformed counts per million (log2CPM) values of all expressed genes (CPM exceeding 4 in at least one group) in our RNA-seq dataset were used for gene ranking. The analysis utilized the h.all.v2023.2.Hs.symbols.gmt Hallmarks gene sets database. GSEA was performed with 1000 permutations, following default settings for the specific comparison outlined in the figure legend.

### Pathway analysis

The pathway analysis focused on investigating terms from Hallmark gene sets (https://www.gsea-msigdb.org/gsea/msigdb/human/collections.jsp#H) within a specific list of genes of interest. Gene sets representing pathways and biological processes were obtained using the R msigdbr package, which provided curated collections of genes associated with specific biological functions and pathways. Overrepresentation Analysis (ORA) was performed using the hypergeometric test to quantify the degree of enrichment, comparing the proportion of genes associated with a particular term within the list to the proportion in the entire genome. Implementation was conducted in R using the clusterProfiler ^81^ package, with subsequent visualization of results through dot plots or bar plots, effectively illustrating significantly enriched terms alongside their corresponding p-values.

### ATAC sequencing

The ATAC library preparation followed a previously reported method ^82^. Briefly, one million cells were lysed using cold lysis buffer (10 mM Tris-HCl, pH 7.4, 10 mM NaCl, 3 mM MgCl2, and 0.1% IGEPAL CA-630), and nuclei were immediately spun at 500x*g* for 10 min in a refrigerated centrifuge. The pellet obtained post nuclei preparation was resuspended in a transposase reaction mix consisting of 25 µl 2× TD buffer, 2.5 µl transposase (Illumina, Cat#: 20034198), and 22.5 µl nuclease-free water. The transposition reaction was carried out for 30 min at 37°C. Following transposition, the sample was purified using a MinElute PCR Purification kit. Library fragments were amplified using 1× NEB next PCR master mix and 1.25 M custom Nextera PCR primers as previously described ^71^, with subsequent purification using a Qiagen PCR cleanup kit, yielding a final library concentration of ∼30 nM in 20 µl. Libraries were amplified for a total of 10–13 cycles and subjected to high-throughput sequencing at the Genomic Resources Core Facility at Weill Cornell Medicine using the Illumina NovaSeq S1 Sequencer with 50-bp paired-end reads. Data from ATAC-seq experiments were derived from four independent experiments with different blood donors.

### ATAC-seq data analysis

For ATACseq data analysis, we utilized the TaRGET-II-ATACseq (https://github.com/Zhang-lab/TaRGET-II-ATACseq-pipeline) pipeline, available on a singularity image (ATAC_IAP_v1.1.simg), to process raw ATACseq data. The read alignments were performed against the GRCh38/hg38 reference human genome. Peak calling was conducted using MACS3 with the following parameters: "macs3 callpeak -f BAMPE -t replicate1 replicate2 replicate3 replicate4 -g hs -q 0.01 --keep-dup 1000 --nomodel --shift 0 --extsize 150". A master consensus peak set was generated by merging the resulting peak files for each treatment condition, followed by merging peaks within 50bp of each other. Quantification of peaks to compare global ATACseq signal changes in the BAM files was conducted using the NCBI/BAMscale program. Raw count matrices were obtained utilizing the BAMscale program. Subsequent analysis utilized the HSS Genomic Core’s reproducible ATACseq analysis pipeline for peak filtering, annotation relative to genomic features, differential peak analysis, and enrichment of signal around specific motifs using ChromVAR. Footprint analyses were performed using TOBIAS according to the user manual. Briefly, we filtered JASPAR2022-CORE_vertebrates transcription factor list based on the TF expressed in our RNAseq data and used this as TF input for motif footprinting. Bioinformatic tools used are listed in Supplementary Table 3.

### Motif Enrichment analysis (HOMER)

*De novo* transcription factor motif analysis was carried out using the motif finder program *findMotifsGenome* in the HOMER ^83^ package, focusing on the given peaks. Peak sequences were compared to random genomic fragments of the same size and normalized G+C content to identify enriched motifs in the targeted sequences.

### Interactive Genome Viewer (IGV)

To visualize the CUT and RUN, and ATAC-seq data, bigwig files were generated for each condition by merging replicate BAM files and then creating normalized coverage bigwig files relative to the sequencing depth using BAMscale ^84^. The normalized bigwig files were then visualized using the IGV browser from the Broad Institute ^85^.

### CUT and RUN

The Epicypher CUT and RUN kit (Cat#: 14-1048) was utilized following the manufacturer’s instructions. To create CUT and RUN libraries, fragmented DNA obtained from the CUT and RUN assay was processed using the NEBNext Ultra II DNA Library Prep Kit for Illumina (NEB, Cat# E7645L), following the manufacturer’s instructions. Libraries were pooled and sequenced (50bp paired end reads) to obtain at least 5 million reads/sample on the Illumina Novaseq SP or Novaseq S4 at the Genomic Resources Core Facility at Weill Cornell Medicine.

### CUT and RUN data analysis

A reproducible CUT and RUN analysis pipeline, CUT&RUNTools 2.0 ^86^ was used to process raw CUT and RUN data. In brief, the sequenced reads were aligned to the human genome (hg38) using bowtie2 ^87^. Peak calling was conducted using MACS2 with the following parameters: "macs2 callpeak - t replicate1 replicate2 replicate3 -g hs -f BAMPE -q 0.01 --scale-to small --nomodel -- keep-dup all". The same approach used for ATAC-seq was applied for downstream analysis. Bioinformatic tools used are listed in Supplementary Table 3.

### Flow cytometric analysis (FACS)

Intracellular flow cytometry was utilized to detect IRF1 and phosphorylated p65 (P-p65) in human monocytes. Prior to cell harvest (5 min) 0.25% Pefablock was added to the cell culture media. Cells were then centrifuged at 300x*g* for 3 minutes, supernatant was discarded, and the cell pellet was gently blotted with a paper towel to remove excess media. Subsequently, cells were washed once with FACS buffer. We used The Foxp3 Transcription Factor Fixation/Permeabilization kit (Invitrogen™, Cat# 00-5521-00) following the manufacturer’s protocol to fix cells for staining with antibodies to transcription factors and nuclear proteins. Briefly, FOXP3 fixation buffer (100 μL per well) was added, and the plate was incubated for 20 minutes at room temperature. After centrifugation, cells were washed again with FACS buffer, followed by resuspension in FACS buffer. Next, cells were incubated with 0.05% Digitonin for 10 minutes at room temperature, followed by centrifugation and a final wash. Fc blocker (BD Biolegend, Cat#:422302) was added to reduce potential nonspecific antibody staining mediated by IgG receptors and incubated for 10 minutes at room temperature. Intracellular antibody staining was performed using an antibody cocktail in 5% BSA-PBS for 60 minutes at room temperature, avoiding light exposure. Finally, cells were washed with FACS buffer and analyzed by flow cytometry. Antibodies and dilutions used are listed in Supplementary Table 2

### CRISPR

For CRISPR-Cas9 genome editing in primary human monocytes, the Lipofectamine™ CRISPRMAX™ Cas9 Transfection Reagent (Invitrogen, Cat#: CMAX00008) was utilized. Initially, cells were resuspended at a concentration of 1.0 x 10^6 cells/mL in media with M-CSF (20ng/mL) and the Jak inhibitor baricitinib (1µM, to prevent activation of Jak-STAT signaling in response to transfected guide RNAs). The cells were then plated at a density of 100,000 cells per well in a 96-well plate and incubated at 37°C for 2-3 hours. Cas9 ribonucleoproteins (Cas9-RNPs) were generated using CRISPRMAX™ Cas9 Transfection Reagent, following the manufacturer’s protocols. Briefly, lyophilized single guide RNAs (sgRNAs) obtained from IDT-DNA (Supplemental Table 1) were reconstituted at a concentration of 50 µM in Nuclease-Free Duplex Buffer (IDT-DNA, Cat# 11-01-03-01). Subsequently, the two sgRNAs per gene were mixed with 61µM Cas9 protein (volume adjusted to achieve equal molar ratio) and incubated with opti-MEM and Cas9 Reagents (part of Lipofectamine kit) for 5 minutes at room temperature to form Cas9-RNPs at a final concentration of 5 µM. For delivery of Cas9-RNP to cells, Lipofectamine CRIPSRMAX lipid nanoparticles were diluted with Opti-mem and incubated for 1min at room temperature. Cas9-RNP and CRISPRMAX mix were incubated at a 1:1 ratio for 15min at room temperature. These Cas9-gRNA-Lipofectamine complexes were added immediately to the plated cells. The cells were then incubated overnight at 37°C. Following transfection, the media was changed to remove residual components, and the cells were further incubated for 24 hours before stimulating them with LPS for 3hours.

### Statistical Analysis

Graphpad Prism for Macs was used for all statistical analysis. Information about the specific tests used, and number of independent experiments is provided in the figure legends. Two-way ANOVA with Tukey’s correction for multiple comparisons was used for grouped data; otherwise, one-way ANOVA with Tukey’s post hoc test for multiple comparisons was performed. Wilcoxon matched-pairs signed rank test was performed for paired non-parametric data.

## Data and code availability

Sequencing data from this study have been deposited at GEO and will be publicly available from the date of publication. Accession numbers are listed in the Supplementary Table. Any additional information required to reanalyze the data (including original code) reported in this paper is available from the first or last authors on request.

## Acknowledgments

We thank the Weill Cornell Medicine Genomics Core for sequencing and the David Z. Rosensweig Genomics Center at HSS for data analysis. The David Z. Rosensweig Genomics Center is supported by The Tow Foundation. Extended Data Fig. 7 was created using Biorender.com.

## AUTHOR CONTRIBUTIONS

B.M. conceptualized, designed, and performed most of the experiments, performed bioinformatic analysis, prepared figures and wrote the manuscript. M.B. performed ATAC-seq experiments. R.Y., V.C., R.B., and C.B., contributed experiments or experimental expertise. L.B.I. conceptualized and oversaw the study and edited the manuscript. All authors reviewed and provided input on the manuscript.

