## Supplemental Figures for "IL-10 Targets IRFs to Suppress IFN and Inflammatory Response Genes by Epigenetic Mechanisms"

### Mishra et al., Extended Data Fig. 1

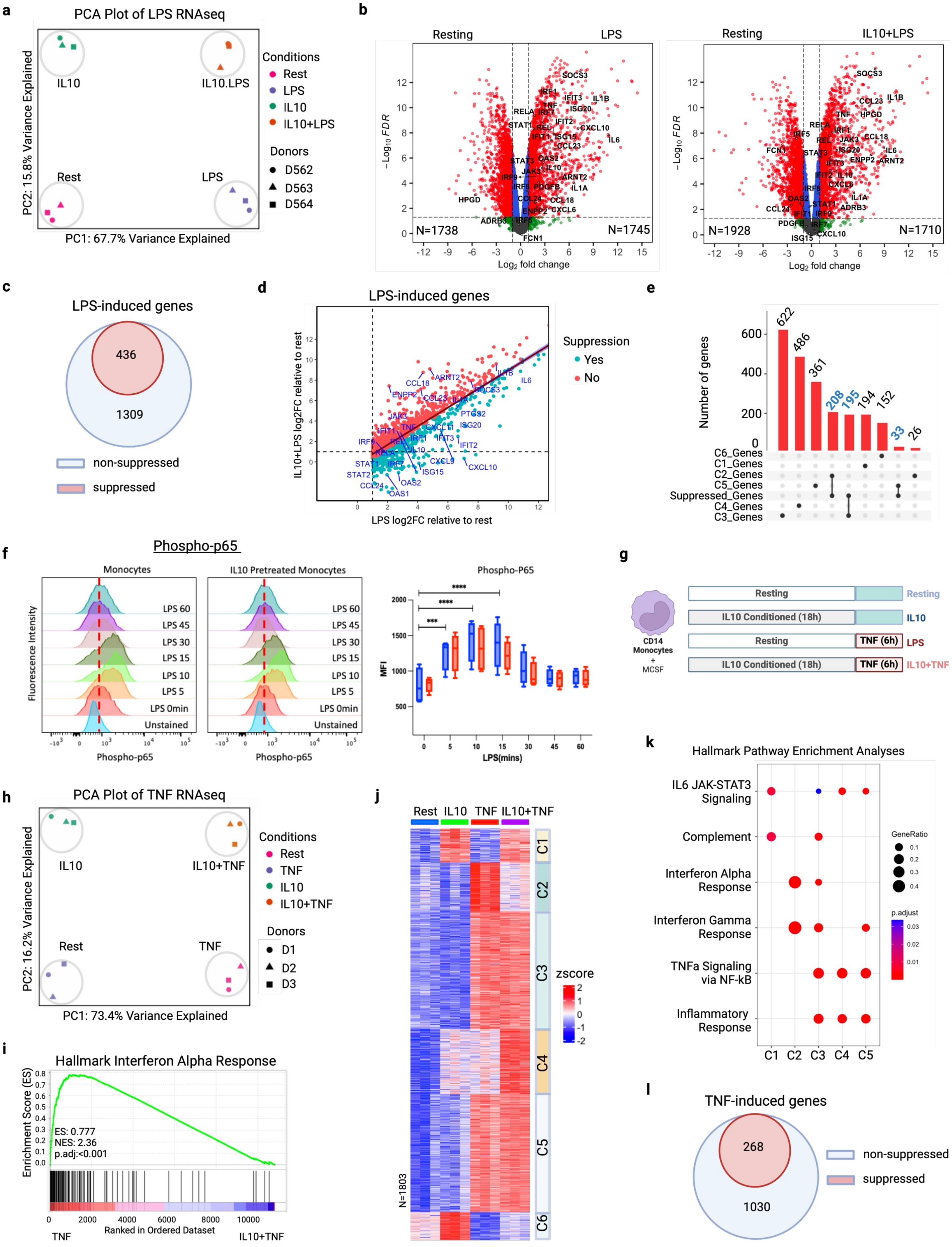

**Mishra et al., Extended Data Fig. 2**

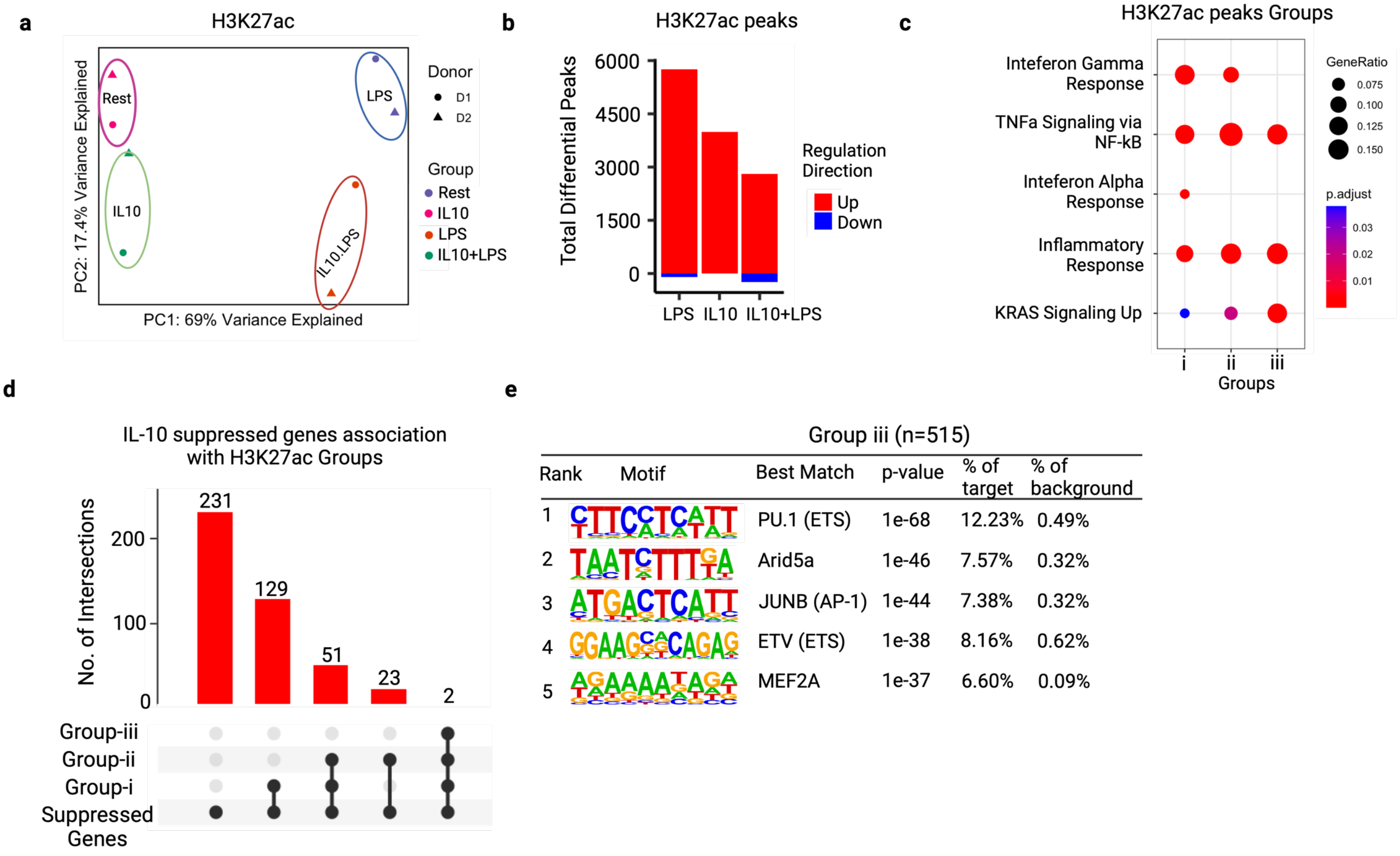

**Mishra et al., Extended Data Fig. 3**

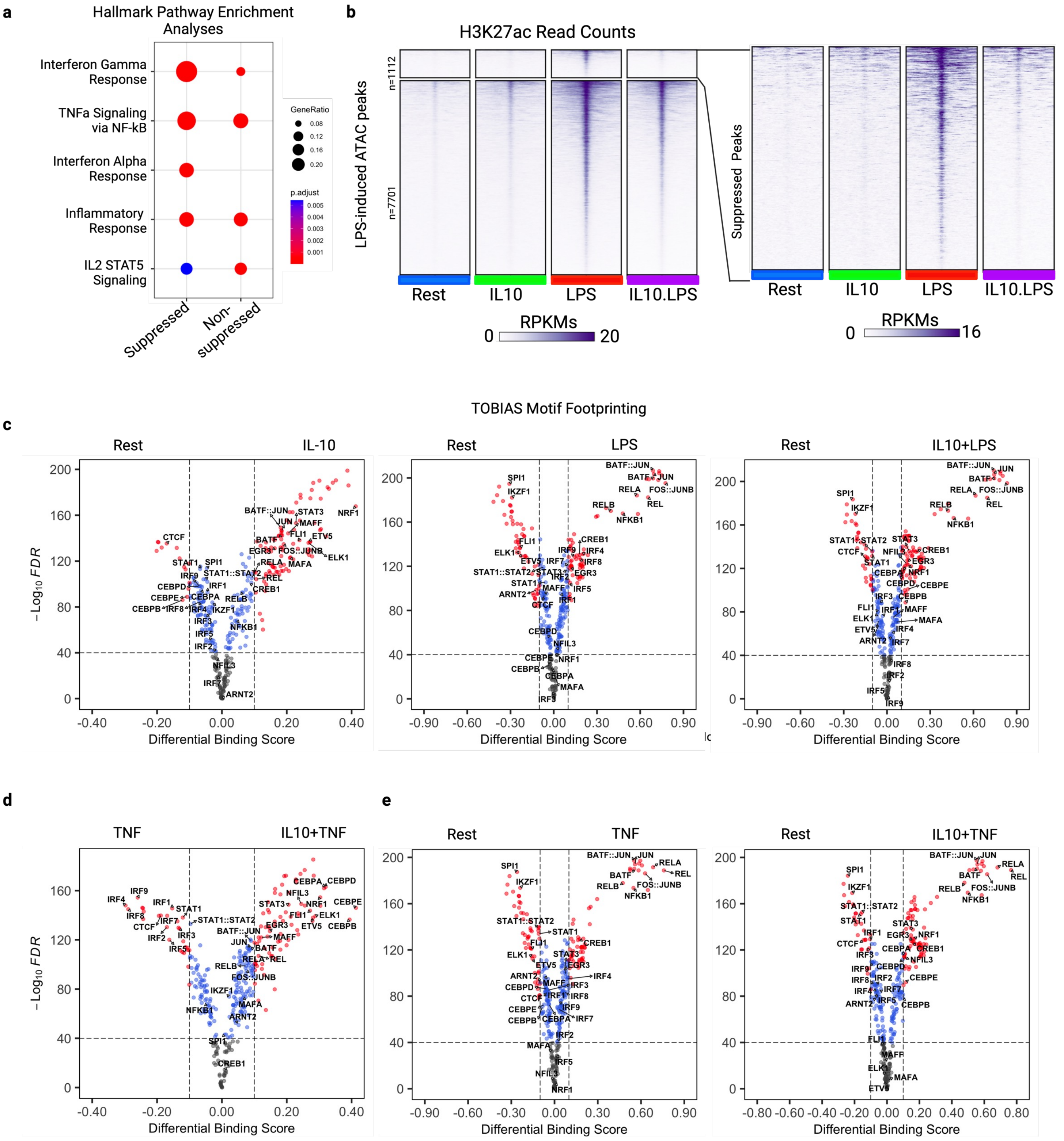

**Mishra et al., Extended Data Fig. 4**

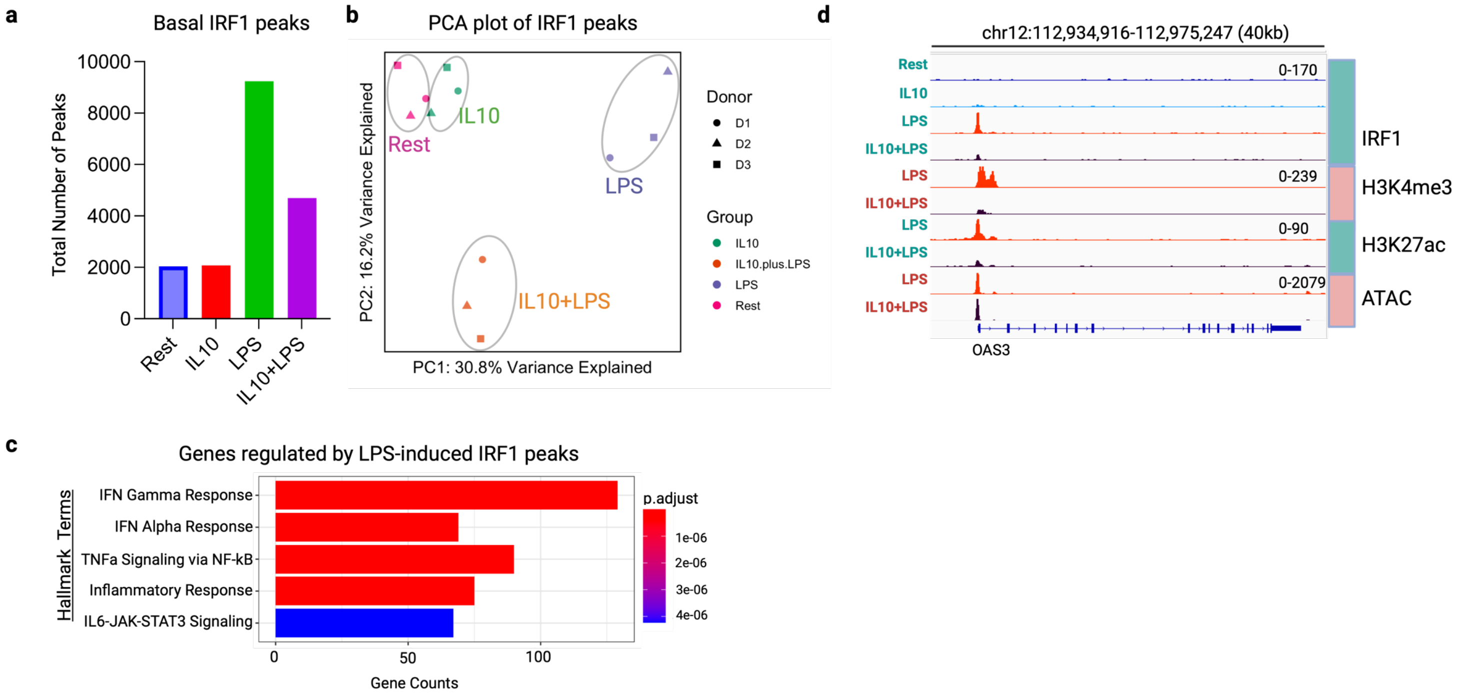

**Mishra et al., Extended Data Fig. 5**

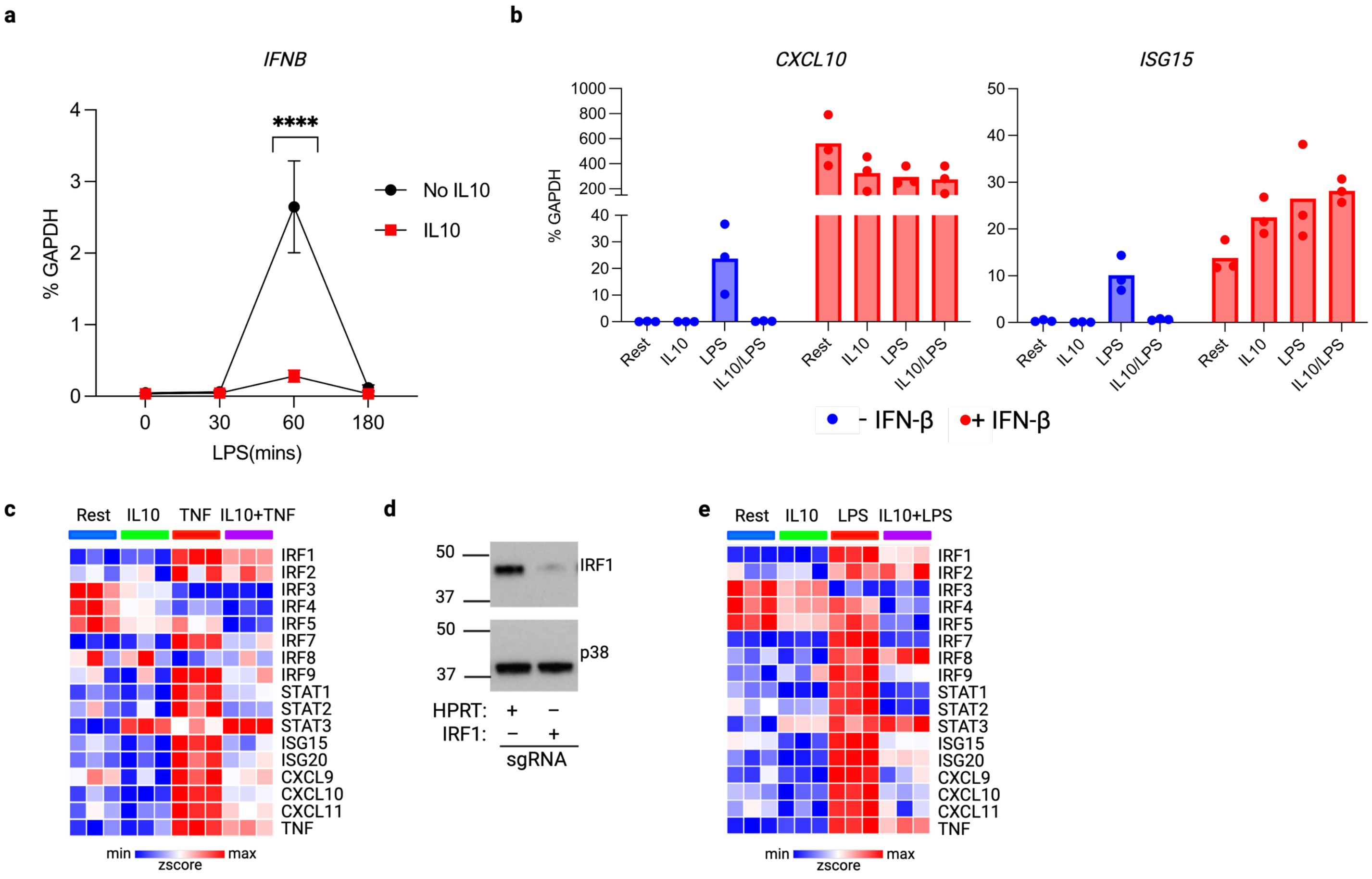

**Mishra et al., Extended Data. 6**

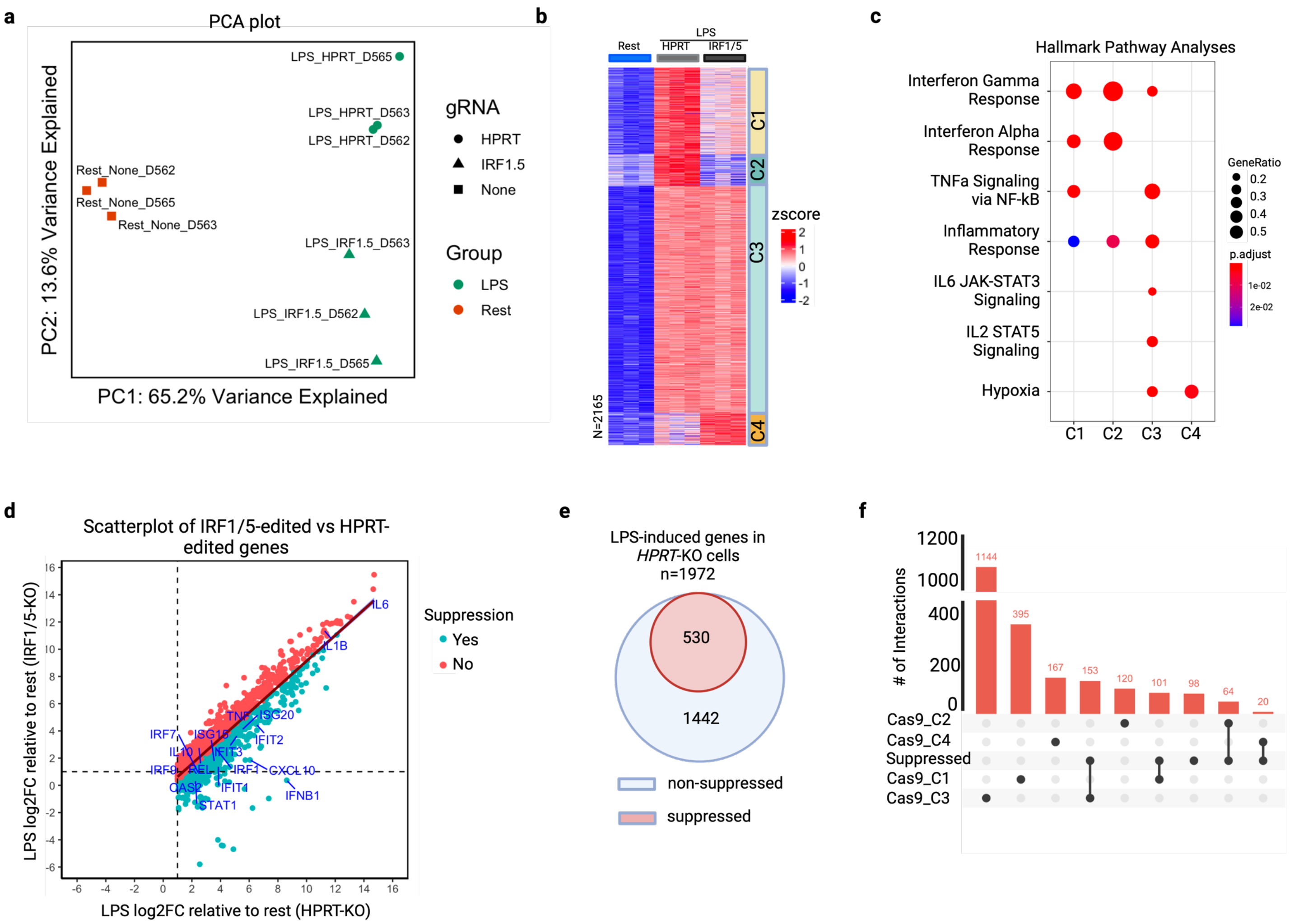

**Mishra et al., Extended Data Fig. 7**

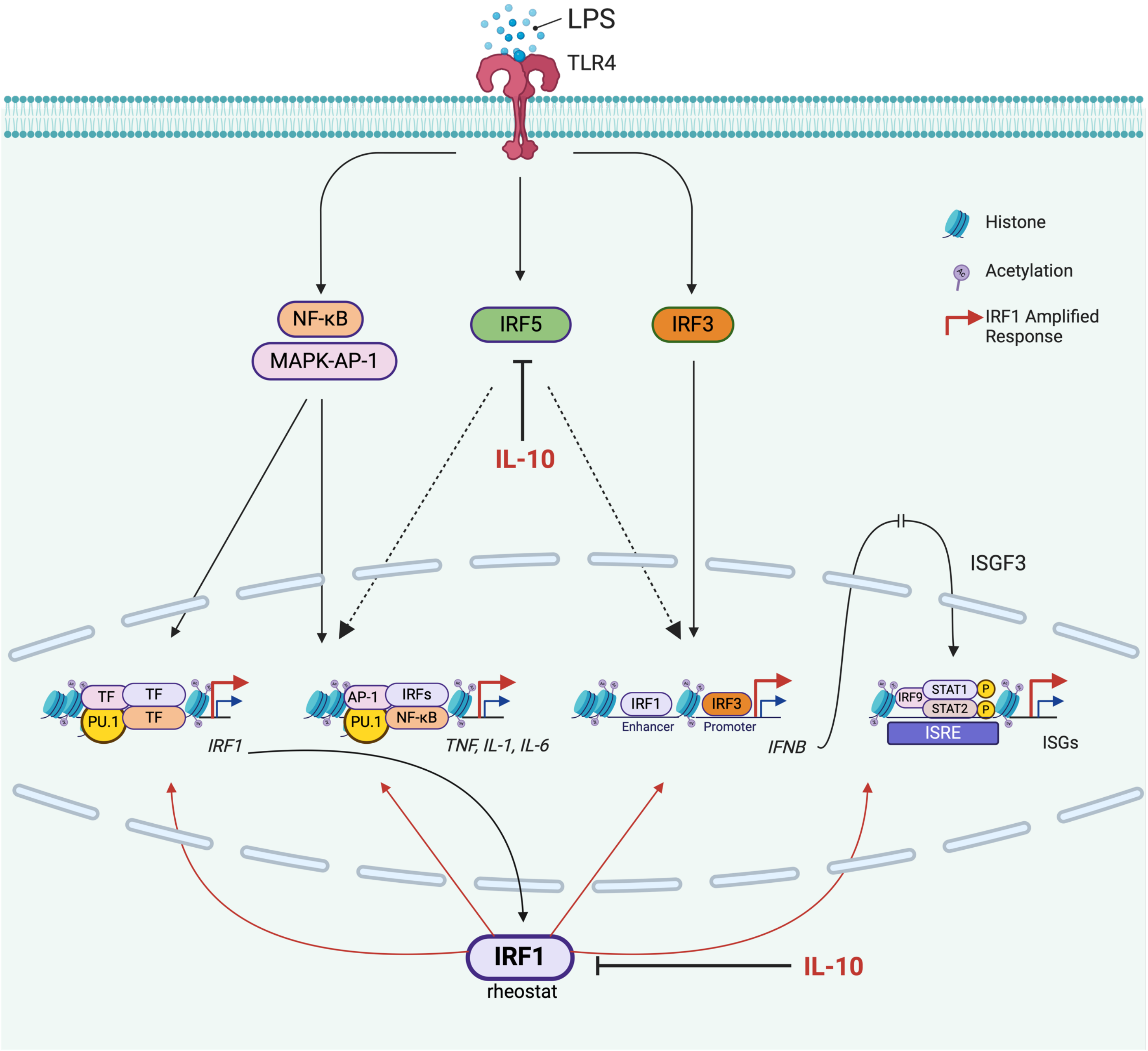
